# Automated slice-specific z-shimming for fMRI of the human spinal cord

**DOI:** 10.1101/2021.07.27.454049

**Authors:** Merve Kaptan, S. Johanna Vannesjo, Toralf Mildner, Ulrike Horn, Ronald Hartley-Davies, Valeria Oliva, Jonathan C.W. Brooks, Nikolaus Weiskopf, Jürgen Finsterbusch, Falk Eippert

**Author notes:** Corresponding authors Address for correspondence: Merve Kaptan & Falk Eippert; Max Planck Research Group Pain Perception, Max Planck Institute for Human Cognitive and Brain Sciences, Stephanstraße 1a, 04103 Leipzig, Germany.

## Abstract

Functional magnetic resonance imaging (fMRI) of the human spinal cord faces many challenges, such as signal loss due to local magnetic field inhomogeneities. This issue can be addressed with slice-specific z-shimming, which compensates for the dephasing effect of the inhomogeneities using a slice-specific gradient pulse. Here, we aim to address outstanding issues regarding this technique by evaluating its effects on several aspects that are directly relevant for spinal fMRI and by developing two automated procedures in order to improve upon the time-consuming and subjective nature of manual selection of z-shims: one procedure finds the z-shim that maximizes signal intensity in each slice of an EPI reference-scan and the other finds the through-slice field inhomogeneity for each EPI-slice in field map data and calculates the required compensation gradient moment. We demonstrate that the beneficial effects of z-shimming are apparent across different echo times, hold true for both the dorsal and ventral horn, and are also apparent in the temporal signal-to-noise ratio (tSNR) of EPI time-series data. Both of our automated approaches were faster than the manual approach, lead to significant improvements in gray matter tSNR compared to no z-shimming and resulted in beneficial effects that were stable across time. While the field-map-based approach performed slightly worse than the manual approach, the EPI-based approach performed as well as the manual one and was furthermore validated on an external corticospinal data-set (N>100). Together, automated z-shimming may improve the data quality of future spinal fMRI studies and lead to increased reproducibility in longitudinal studies.

## 1. Introduction

The spinal cord is one of the key structures linking the brain with the peripheral nervous system and participates in numerous sensory, motor and autonomic functions (Hochman, 2007). Non-invasive approaches to investigate the human spinal cord are therefore of great interest not only from a basic neuroscientific perspective, but also with regards to their possible clinical utility in order to understand pathological mechanisms in motor and sensory disorders such as multiple sclerosis and chronic pain (Wheeler-Kingshott et al., 2014). Currently, the main approach to investigate spinal cord function is based on blood-oxygen-level-dependent functional magnetic resonance imaging (BOLD fMRI; for reviews see Giove et al., 2004; Stroman et al., 2014; Summers & Brooks, 2014; Cohen-Adad, 2017). Using conventional BOLD fMRI techniques such as gradient-echo echo-planar imaging (GE EPI) is however challenging in the spinal cord due to i) its small cross-sectional diameter, ii) prominent physiological noise from cardiac and respiratory sources, and iii) magnetic field inhomogeneities.

In the cervical spinal cord (i.e. the part that is easiest to access with currently available receive coils at 3T), inhomogeneities in the magnetic field occur at both large and small spatial scales (Cohen-Adad, 2017). While large-scale variations are for example due to the proximity of the lungs (and can thus vary dynamically; e.g. Verma & Cohen-Adad, 2014), small-scale variations are due to the interfaces between vertebrae and connective tissue, which have different magnetic susceptibilities (Cooke et al., 2004; Finsterbusch et al., 2012). These small-scale field inhomogeneities are reproduced spatially along the superior-inferior axis of the spinal cord and significantly affect image quality, leading to consistent patterns of signal loss (Maieron et al., 2007; Finsterbusch et al., 2012). While it would thus be imperative for reliable and reproducible fMRI of the spinal cord to mitigate these effects, standard shimming techniques implemented on common whole-body MR systems are not able to compensate these spatially repeating inhomogeneities to an adequate degree (Finsterbusch, 2014).

One method that is commonly employed to overcome through-slice dephasing is slice-specific ’z-shimming’ (Frahm et al., 1988; Constable, 1995; Glover, 1999) where an additional gradient pulse is applied in the slice-selection direction in order to compensate the effect of susceptibility-induced gradients and resulting signal loss. In the brain, z-shimming has been applied in GE EPI studies focused on susceptibility-prone regions, i.e. those that are close to air/bone interfaces such as the orbitofrontal, the medial temporal, and the inferior temporal lobes (Yang et al., 1997; Deichmann et al., 2003; Posse et al., 2003; Weiskopf et al., 2006). Finsterbusch et al. (2012) investigated whether one could use this approach to also compensate for the periodically occurring signal drop- outs (along the superior-inferior axis) on T2*-weighted GE EPI images of the spinal cord. By applying *single*, slice-specific compensation moments – which were manually determined based on a reference-scan acquired prior to the experimental EPI acquisition – they were able to demonstrate an improvement in spinal cord image quality: reducing the spatially repeating signal drop-outs via slice-specific z-shimming resulted in an increase of mean signal-intensity by ∼20% and a reduction of signal-intensity variability along the cord by ∼80%.

While the slice-specific z-shimming protocol developed by Finsterbusch and colleagues has already been used in numerous spinal (e.g. Sprenger et al., 2012; Geuter & Buchel, 2013; Kong et al., 2014; van de Sand et al., 2015; Eippert et al., 2017; Sprenger et al., 2018) and cortico-spinal fMRI studies (e.g. Sprenger et al., 2015; Tinnermann et al., 2017; Vahdat et al., 2020; Oliva et al., 2022), the impact of slice-specific z-shimming on EPI time-series data has not been investigated systematically, as Finsterbusch and colleagues only evaluated its effects on single volumes of GE EPI data, but not on time-series metrics such as tSNR (Welvaert & Rosseel, 2013). Even more important – and already argued for by Finsterbusch and colleagues – would be an automated way to determine the slice-specific z-shims, as these are currently determined manually by the scanner operator: either visually by going through each slice and z-shim value obtained in a reference-scan or by manually placing a region of interest on each slice of this reference scan and evaluating the extracted signal intensity. This procedure is time-consuming, requires expertise in judging the quality of spinal EPI data, and contains a subjective component, thus also limiting its potential in terms of reproducibility.

In this study, we aim to develop an automated and user-friendly procedure for determining slice-specific z-shims in order to improve the quality of spinal fMRI. In a first step, we aim to replicate the results of Finsterbusch et al. using twice the original sample size (N=48). Next, we aim to extend their findings by probing the relevance of slice-specific z-shimming for fMRI through investigating its effects a) across different echo times, b) in distinct anatomical regions, and c) on a time-series metric (tSNR). Most importantly, we propose two different automated methods for determining slice-specific z-shims (each based on a sample size of N=24). The first method is based on a z-shim reference-scan acquisition and determines z-shim values by analyzing EPI signal intensity within the spinal cord for each combination of slice and z-shim value. The second method is based on a field map acquisition and determines z-shim values by estimating the strength of the gradient field needed to compensate for the local through-slice inhomogeneity for each slice. In a final step, we use an independently-acquired external data-set (N>100; Oliva et al., 2022) in order to validate our candidate approach for automating the selection of slice-specific z-shims.

## 2. Material and Methods

### 2.1 Participants

48 healthy participants (22 females, mean age: 27.17 years, range 20-37 years) participated in this study. All participants provided written informed consent and the study was approved by the ethics committee at the Medical Faculty of the University of Leipzig. The sample size was determined based on a study by Finsterbusch et al. (2012): as we wanted to replicate and extend their findings (which were based on a sample of N=24), we chose the same sample size for each of our two sub- groups, resulting in an overall sample size of N=48.

### 2.2 Study design

All participants underwent the following scans in the order described below (for details of scans, see section ‘2.3 Data acquisition’).

After an initial localizer scan, the EPI slice stack and the adjust volume were prescribed and a single EPI volume was acquired in order to initialize the scanner’s ‘Advanced shim mode’ – this shim was then employed in all the following EPI acquisitions by using the same adjust volume. An EPI z-shim reference scan was performed next in order to allow for the manual as well as EPI-based automated selection of the optimal z-shim moment for each slice. Two sagittal field maps (vendor-based and in-house versions, respectively) were then acquired to obtain the B_0_ static magnetic field distribution, of which the vendor-based one was used for the field map based automated z-shim selection due to it being widely available. This was followed by the acquisition of a high-resolution T2-weighted image in order to allow for spinal cord segmentation as needed for the field map based automated z-shim selection.

In order to compare the signal characteristics under different z-shimming conditions, EPI data were acquired with three different EPI protocols for each participant: without z-shim gradient compensation (condition “no z-shim”), with z-shim gradient compensation based on manual z-shim selection (condition “manual z-shim”), and with z-shim gradient compensation based on automated z-shim selection (condition “automated z-shim”). For one-half of the participants (24 participants), the automated selection was based on the EPI reference scan, whereas for the other half, the automated selection was based on the vendor-based field map. Both single EPI volumes (as in Finsterbusch et al., 2012), as well as 250 EPI volumes (in order to assess effects on time-series data), were acquired for each condition; the order of the EPI scans under different conditions was pseudo-randomized across participants.

We also wanted to assess the benefits of slice-specific z-shimming at different echo times (TE), and therefore acquired 25 EPI volumes under three different TEs (30, 40, and 50ms, each with a repetition time (TR) of 2552ms) for each of the three conditions (please note that the z-shim indices chosen reflect gradient fields to be compensated – rather than moments of the compensation gradient pulse – and thus scale the pulsed gradient moment with the TE such that a determined index is valid for all TEs). The order of the EPI scans acquired with different TEs were also pseudo-randomized across participants.

The EPI reference scan and the in-house field map acquisitions were repeated at the end of the scanning session in order to assess the stability of z-shimming across time.

### 2.3 Data acquisition

All measurements were performed on a 3T whole-body Siemens Prisma MRI System (Siemens, Erlangen, Germany) equipped with a whole-body radio-frequency (RF) transmit coil and 64- channel RF head-and-neck coil and a 32-channel RF spine-array, using the head coil element groups 5–7, the neck coil element groups 1 and 2, and spine coil element group 1 (all receive-only).

EPI acquisitions were based on the z-shim protocol developed by Finsterbusch et al. (2012) that employed a *single*, slice-specific gradient pulse for compensating through-slice signal dephasing. EPI volumes covered the spinal cord from the 2nd cervical vertebra to the 1st thoracic vertebra and were acquired with the following parameters: slice orientation: transverse oblique; number of slices: 24; slice thickness: 5mm; field of view: 128×128mm^2^, in-plane resolution: 1×1mm^2^; TR: 2312ms; TE: 40ms; flip angle: 84°; GRAPPA acceleration factor: 2; partial Fourier factor: 7/8, phase-encoding direction: anterior-to-posterior (AP), echo spacing: 0.93ms, bandwidth per pixel: 1220 Hz/Pixel; additionally, fat saturation was employed. The EPI reference scan (TE: 40ms, total acquisition time: 55 seconds) was acquired with 21 equidistant z-shim moments compensating field inhomogeneities between +21 and -21 mT m^-1^ms (in steps of 2.1 mT m^-1^ms).

The vendor-based field map (total acquisition time: 4.31min) was obtained using the 2D GRE sequence provided by Siemens with two echoes per shot (TE 1: 4.00ms; TE 2: 6.46ms; slice orientation: sagittal (parallel to the normal vector of the axial EPI slices); slice number: 32; slice thickness: 2.2mm; field-of-view: 180×180mm^2^; in-plane resolution: 1×1mm^2^; TR: 500ms; flip angle: 50°, bandwidth per pixel of 1030 Hz/pixel). Additionally, an in-house field map based on a 3D multi-echo FLASH sequence with multiple gradient echoes acquired at short inter-TEs was acquired, which yielded a superior signal-to-noise ratio at a reduced overall scan time. This contained 12 bipolar gradient echoes (which allowed for shorter inter-echo spacings; note that potential image shifts were avoided by a multi-echo navigator scan without phase encoding right at the start of image acquisition; a phase correction between the odd and even echoes was performed by the vendor’s Ice reconstruction pipeline), a TE increment/difference of 1.3ms, fat suppression RF pulses with corresponding spoiler gradients before each slab-selective excitation, a repetition time of 32ms, a flip angle of 15°, bandwidth per pixel of 1030 Hz/pixel, and sagittal slice orientation (parallel to the normal vector of the axial EPI slices). The in-plane and partition resolutions of this in-house field map were 1×1mm^2^ and 2.2mm, respectively, with corresponding fields-of-view of 180×180×70.4mm^3^. A total scan time of less than 2min was achieved by the application of GRAPPA (an acceleration factor of 2 was used in PE dimension). The frequency offset Δ*v*_0_ in each voxel was extracted from a linear fit to the unwrapped phases of all echoes (unwrapping of phase jumps exceeding +/- Pi was performed using a simple algorithm; due to the employed short echo and inter-echo times, this unwrapping could be applied because problems of noisy phase jumps or an undersampling of the phase evolution were largely absent).

A high-resolution T2-weighted image was acquired using a 3D sagittal SPACE sequence as recently recommended (Cohen-Adad et al., 2021; 64 sagittal slices; resolution: 0.8×0.8×0.8mm^3^; field-of-view 256×256mm^2^; TE: 120ms; flip angle: 120°; TR: 1500ms; GRAPPA acceleration factor: 3; acquisition time: 4.02min).

### 2.4 Selection of slice-specific z-shim moments

#### 2.4.1 Manual selection

The researcher carrying out the data acquisition (MK) determined the z-shim moment with the highest signal intensity in the spinal cord for each slice by visual inspection (i.e. for each of the 24 slices, the researcher looked at all 21 volumes – each volume reflecting an acquisition with one z-shim moment – in order to determine the “optimal” z-shim moment for each slice). This selection process took ∼10 minutes per participant and was carried out for all 48 participants, i.e. in both sub-groups of 24 participants.

#### 2.4.2 Automated selection

The necessary scans for the automated selection (EPI reference-scan for EPI-based selection; vendor-based field map and T2-weighted scan for field map based selection) were sent from the scanner console to the online calculation computer (OS: Ubuntu 18.04, CPU: Intel Core(TM) i7- 3770K 3.50GHz, RAM: 16 GB, Mainboard: Gigabyte Z77X-UD3H) using the scanner console’s in-built network connection. In-house MATLAB (The Mathworks Inc, 2019) scripts utilizing tools from dcm2niix (version 1.0.20180622; Li et al., 2016; https://github.com/rordenlab/dcm2niix), SCT (version 3.2.7; De Leener et al., 2017; https://spinalcordtoolbox.com/en/stable/), and FSL (version 5.0; Jenkinson et al., 2012; https://fsl.fmrib.ox.ac.uk/fsl/fslwiki) were employed to determine the optimal z-shim moment for each slice. These values were then sent back to the scanner console in a text file that is read by the z-shim sequence. An overview of the automated methods is given in Figure 1 (please note that the z-shim selection process is automated and does not require any input from the user).

**Figure 1.**
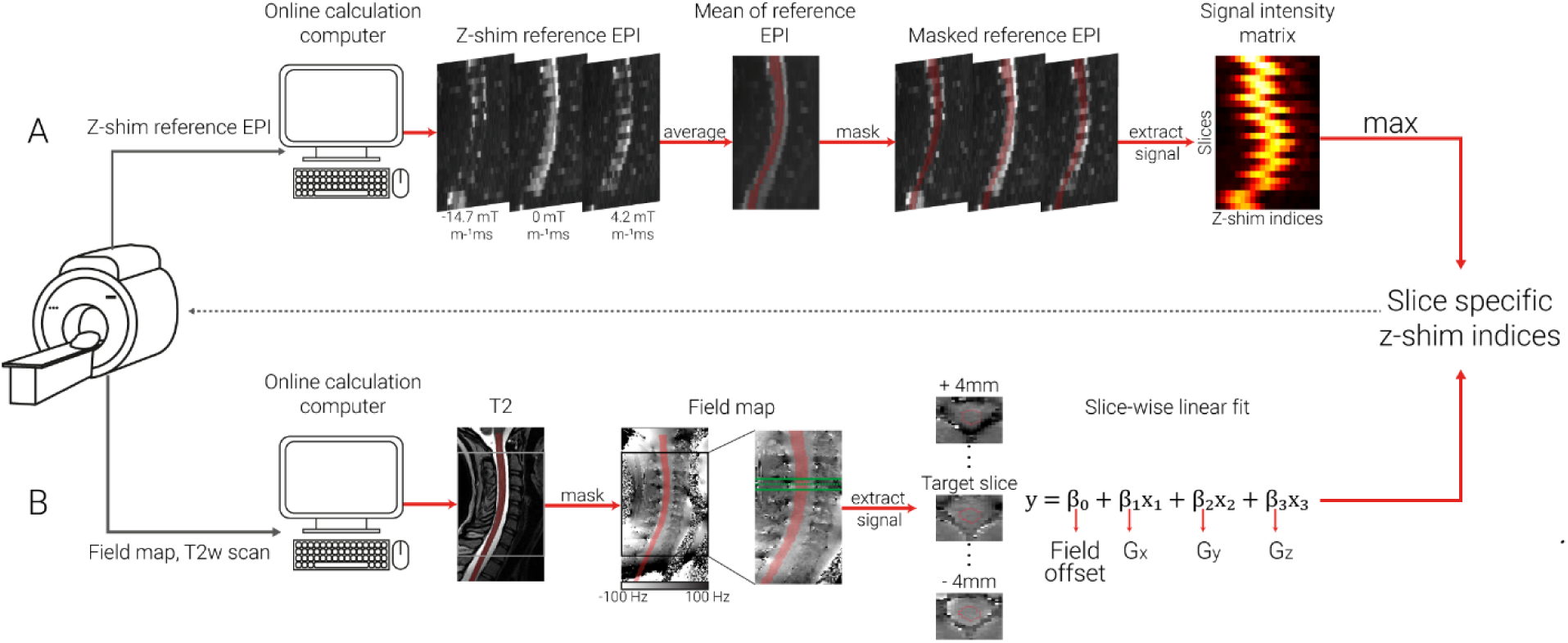
Schematic depiction of automated z-shim methods. After the acquisition of the necessary scans for each method (z-shim reference EPI for EPI-based approach, T2-weighted image and field map for field map based approach), DICOM images were exported to an online calculation computer, and converted to NIfTI format before further processing. **A. EPI-based selection**. The z-shim reference scan was then averaged across its 21 volumes (one volume per z-shim moment; three volumes are depicted here as mid-sagittal sections to illustrate the varying signal loss) and the resulting mean image was segmented (the segmentation is shown here as a transparent red overlay for display purposes). The mean signal intensities for each slice and z-shim moment were extracted from the segmented cord, resulting in a 24×21 signal intensity matrix (slices×volumes). For each slice, the z-shim value (i.e., the corresponding index in the reference scan) resulting in the maximum intensity was selected. **B. Field map based selection**. A high-resolution T2-weighted image was segmented and used to determine the field map voxels to be included in the fitting procedure (the segmentation is shown as a transparent red overlay for display purposes). The gray and the black boxes depict the EPI coverage on the T2-weighted image and phase map, respectively. Voxels within a 9mm thick slab (i.e. 9 transversal field map slices, corresponding to a 5mm EPI slice + 2mm on each side) were included in a slice-wise fitting procedure. The green lines on the phase map indicate the input volume for fitting an exemplary target slice (dashed green line). Exemplary transversal slices are also shown, with the red line outlining the spinal cord. Slice-wise fitting, including three linear field coefficients (G_x_, Gy and G_z_) along the main axes of the imaging volume and a spatially homogenous field term (field offset), was repeated over slices and the z-shim (G_z_) moments corresponding to the center ofthe EPI slices were selected.

##### 2.4.2.1 EPI-based selection

In a subsample of 24 participants, the EPI z-shim reference-scan was used to determine the optimum z-shim moments. The EPI z-shim reference-scan – consisting of 21 volumes (each volume corresponding to one z-shim moment) with 24 slices each – was then averaged over the 21 volumes (i.e. over all z-shim moments) and the resulting mean image was automatically segmented using the PropSeg approach implemented in SCT (De Leener et al., 2014). Based on experience from pilot experiments, we built in several fail-safes (i.e. systematically changing the arguments of SCT’s PropSeg function that affect the propagation in the z-direction) in order to ensure that the segmentation would propagate across the entire slice stack; this possibility to automatically adjust parameters in case of failure was also the reason that – out of SCT’s segmentation algorithms – we chose PropSeg instead of DeepSeg. We used the mean image for segmentation because we wanted to ensure that image quality was sufficient for automatic segmentation of the spinal cord and because the averaging of volumes acquired during different breathing cycles avoids a bias towards one respiratory state as could occur with single volumes. In post-hoc investigations regarding the suitability of using the mean EPI image for segmentation, we i) used a maximum image instead of a mean image as the input for segmentation and ii) used a segmentation obtained from the T2-weighted image (registered to the EPI segmentation), but both of these alternative approaches resulted in highly similar results compared to our original approach (data not shown). Using the automatically generated spinal cord mask, the mean signal intensities for each slice and z-shim moment were extracted, resulting in a 24×21 matrix, from which the z-shim moment yielding the maximum intensity across the cord mask was determined for each slice. The average run-time for the execution of the selection code was 15.6 seconds (range across the entire sample: 7.7- 62.3 seconds), with the variation mostly being due to the number of PropSeg runs needed to achieve complete propagation. The interested reader can assess the quality of the EPI-based spinal cord segmentation via a quality-control HTML-report shared together with our data-set (see section 2.8).

##### 2.4.2.2 Field map (FM) based selection

In another subsample of 24 participants, sagittal field maps (acquired with the same angulation as EPI data) were used to determine the optimum z-shim moments; note that field maps had anisotropic voxels, as i) a high in-plane resolution of the sagittal field map is necessary in order to obtain sufficient information about the gradient in the through-slice direction of the EPI (i.e. foot-head) and ii) the left-right direction (where voxels were largest) is expected to have the least field variation and is thus least sensitive to resolution. First, a spinal cord mask was generated via a PropSeg-based automatic segmentation of each participant’s T2-weighted image because a high-quality segmentation of the field map magnitude image was not possible due to the sagittal slice thickness of 2.2mm as well as the poor image contrast between spinal cord and cerebrospinal fluid (note that since the T2-weighted image and field map were well aligned and acquired right after each other we did not carry out a separate registration step). Field map based (from now on referred to as FM-based) z-shim moments were then calculated for each EPI slice using a linear least-squares fit of a set of spatial basis functions to the measured field map (which was smoothed with an isotropic 1mm Gaussian kernel prior to the calculation). The spatial basis functions consisted of three linear field terms along the main imaging axes and a spatially homogenous field term, representing a field offset (although obtaining x- and y-gradients is not necessary for calculating the through-slice field component, their inclusion can be seen as a step towards full slice-wise shimming [see also Islam et al., 2019] and obtaining y-gradients is necessary for determining the effective TE [see below]). Only voxels within the spinal cord mask contributed to the fitting procedure, which included voxels within a 9mm thick slab (i.e. 9 transversal field map slices) centered on the center of the corresponding EPI slice. The slab was chosen to be thicker than the EPI slice (i.e. an additional 2mm either side) in order to give more robust estimates of the through-slice field gradient. The fitted through-slice linear field term (*G_z_*) was taken to represent the local field gradient causing through-slice signal dephasing within the corresponding EPI slice. The resulting dephasing gradient moment of *G_z_* · *TE* was rounded to the nearest of the 21 z-shim compensations available in the EPI protocol and then used for subsequent EPI acquisitions. The average time for the execution of the selection code was 36.1 seconds (range across the entire sample: 31.5- 53.3 seconds).

### 2.5 Preprocessing

All images were visually inspected before the analysis for potential artefacts. Preprocessing steps were performed using MATLAB (version 2021a), FSL (version 6.0.3), and SCT (version 4.2.2; please note that a more recent version of SCT was used for preprocessing (4.2.2) compared to the automated analysis during data acquisition (3.2.7), due to the availability of releases at the respective times). The reason we carried out preprocessing steps and did not work only on the raw data is two-fold: i) we were interested in z-shim effects on time-series metrics (tSNR) and thus needed to motion-correct the EPI time-series data and ii) we were performing most analyses in template space and thus need to bring structural and functional data to this space (requiring segmentation and registration-to-template steps). Please note that – depending on context – we are using the terms “fMRI data” and “EPI time-series data” interchangeably.

#### 2.5.1 Motion-correction of EPI time-series data

A two-step motion correction procedure (with spline interpolation) was applied to the EPI time-series data. Initially, the mean of 750 volumes (250 volumes under each of the three different conditions, i.e. no z-shim | manual z-shim | automated z-shim) was calculated in order to serve as the target image for the first step of motion correction; averaging across all three conditions eliminates a bias towards any one condition with respect to the target image. Based on this mean image, the spinal cord was automatically segmented in order to provide a spinal cord centerline that then served as input for creating a cylindrical mask (with a diameter of 30mm). This mask was employed during the motion-correction procedure in order to ensure that image regions moving independently from the cord would not adversely affect motion estimation. Slice-wise motion correction with a 2^nd^ degree polynomial regularization in the z-direction was then performed (De Leener et al., 2017). In the second step, a new target image was obtained by calculating the mean of motion-corrected images from the first step and the raw images were realigned to this new target image, using the identical procedure as described above. Please note that the data obtained under different TEs (25 images per TE and condition) were also registered to this target image using the same procedure.

Under the “no z-shim” condition, especially the inferior slices suffered from severe signal drop- outs that hampered the quality of the slice-wise motion correction algorithm by inducing ’artificial’ movements that were indeed not present in the raw data. This could impact the tSNR calculation negatively by artificially increasing the standard deviation across time and thus give an inflated estimate of the beneficial effects of z-shimming. Therefore, in a control analysis, we also performed a ‘censoring’ of outlier volumes before the tSNR calculation. The outlier volumes were defined using dVARS (the root mean square difference between successive volumes) and refRMS (root mean square intensity difference of each volume to the reference volume) as metrics using FSL’s ‘fsl_motion_outliers’ tool. Volumes presenting with dVARS or refRMS values two standard deviations above the mean values of each run were selected as outliers. These outlier volumes were then individually modelled as regressors of no interest.

#### 2.5.2 Segmentation

T2-weighted images were initially segmented using the DeepSeg approach implemented in SCT (Gros et al., 2019). This initial segmentation was used for smoothing the cord along its centerline using an anisotropic kernel with 8mm sigma. The smoothed image was again segmented in order to improve the robustness of segmentation. The quality of the segmentations was assessed visually and further manual corrections were not deemed to be necessary in any participant.

For functional images, a manual segmentation was used instead of an automated procedure, as the registration to template space relied on segmentations and we therefore aimed to make this preprocessing step as accurate as possible. For the single-volume EPIs, the single volumes under the three different z-shimming conditions were averaged and this across-condition mean image was used to manually draw a spinal cord mask. For the EPI time-series, all motion-corrected volumes were averaged and a spinal cord mask was manually drawn based on this mean image (please note that this mask was also used for the normalization of the volumes with different TEs). These manually drawn masks were also used to calculate results in native space.

#### 2.5.3 Registration to template space

SCT was utilized for registering the EPI images to the PAM50 template space (De Leener et al., 2018); PAM50 is an MRI template of the spinal cord and brainstem available in SCT for multiple MRI contrasts. The T2-weighted image of each participant was brought into template space using three consecutive registration steps: i) using the spinal cord segmentation, the spinal cord was straightened, ii) the automatically determined labels of vertebrae between C2-C7 (manually corrected where necessary) were used for vertebral alignment between the template and the individual T2-weighted image, and iii) the T2-weighted image was registered to the template using non-rigid segmentation-based transformations.

In order to bring the functional images to template space, the template was registered to the functional images using non-rigid transformations (with the initial step using the inverse warping field obtained from the registration of the T2-weighted image to the template image). The resulting inverse warping fields obtained from this registration (from native EPI space to template space) were then applied to the respective functional images (e.g. single EPI volumes, mean EPI volume, tSNR maps) to bring them into template space where statistical analyses were carried out.

Finally, we also brought each participant’s field map into template space in order to visualize the average B_0_ field variation across participants. Each participant’s field map was first resampled to the resolution of the T2-weighted image before the warping field obtained from the registration to template space was applied to the field map.

#### 2.5.4 EPI signal extraction

In order to assess the effects of z-shimming, we obtained signal intensity data from each EPI slice. When analyses were carried out in native space and were based on the entire spinal cord cross-section, we used the above-mentioned hand-drawn masks of the spinal cord and obtained one value per slice (average across the entire slice). In contrast, when analyses were carried out in template space or were based on gray matter regions only, we made use of the available PAM50 template masks of the entire spinal cord or the gray matter (with the probabilistic gray matter masks thresholded at 90%); again, we obtained one average value per mask and slice. Please note that in addition to reporting p-values from statistical tests, we also report (where appropriate) the percentage difference between conditions and the associated 95% confidence interval (CI) as estimated via bootstrapping.

### 2.6 Statistical analysis

#### 2.6.1 Replication and extension of previous findings

##### 2.6.1.1 Direct replication

In a first set of analyses (across all 48 participants), we aimed to replicate the findings of Finsterbusch et al. (2012). We, therefore, used template space single-volume EPI data acquired under no z-shim and manual z-shim conditions, calculated the individual EPI signal intensity per slice and reported the *mean* of signal intensity across all slices as well as the *variation* of signal intensity across all slices; for the latter, we initially used the *variance* (as done by Finsterbusch et al., 2012), but after the replication of their results we employed the *coefficient of variation* for the remainder of the manuscript (due to it being a standardized measure of variability). Both descriptive changes (percent increase / decrease), as well as statistical values (based on paired t-tests), were reported for the condition comparison. To additionally investigate how robust these findings were, we complemented these single-volume analyses – that might be affected by various noise sources – by the same analysis approach, but now carried out on an EPI volume that is the average of a time-series of 250 motion-corrected EPI volumes (acquired both for no z-shim and manual z-shim; Supplementary Material). In order to demonstrate that neither of these results were impacted by registration to template space, we also reported native space results in the Supplementary Material.

##### 2.6.1.2 Slice-by-slice characterization of z-shim effects

Finsterbusch et al. (2012) already demonstrated that the improvement due to slice-specific z-shimming varies spatially along the rostro-caudal direction. We therefore reasoned that it might be informative to also quantify the benefit for slices with various degrees of signal-loss (obviously, such an analysis could only be performed in native space). We first did this in a descriptive manner by reporting i) the maximally found percentage increase in signal intensity due to z-shimming and ii) the proportion of slices that differed by 0, 1, 2, 3, and >3 z-shim steps from the ‘neutral’ setting of no z-shim. In the Supplementary Material, we then followed this up more formally with an analysis where we categorized slices according to the manually chosen z-shim value and compared the signal intensity in these categories between no z-shim and manual z-shim both descriptively (using % signal intensity difference) and inferentially using a 2×5 repeated-measures ANOVA (factor 1: condition with two levels: no z-shim, manual z-shim; factor 2: step-difference with five levels: 0, 1, 2, 3, >3). We tested for a main effect of condition, a main effect of step-difference and an interaction between these two factors; post-hoc t-tests were Bonferroni corrected. To estimate the robustness of the results from these analyses (which were based on single EPI volumes), we repeated them on the average across the 250 motion-corrected EPI volumes (Supplementary Material).

##### 2.6.1.3 z-shim effects across different TEs

We also aimed to assess the effects of z-shimming at TEs clearly shorter (30ms; fastest TE possible with the employed partial-Fourier factor of 7/8) and longer (50ms; same distance to our standard TE of 40ms) than the estimated T2* in the cervical spinal cord at 3T (∼40ms; Barry et al., 2019), considering that such choices might often be necessary in fMRI studies. We, therefore repeated the analyses described in section 2.6.1.1 (assessing the *mean* of signal intensity across all slices as well as the *variation* of signal intensity across all slices for no z-shim and manual z-shim conditions) on the template-space EPI data obtained with TEs of 30ms and 50ms, both for single-volume data and (in the Supplementary Material) for an average of the 25 volumes acquired at each of the different TEs.

##### 2.6.1.4 z-shim effects in gray matter regions

The effects reported in Finsterbusch et al. (2012) were obtained from averages across the entire cross-section of the spinal cord, thus mixing gray and white matter signals. However, with the availability of probabilistic gray matter maps (via SCT, see https://github.com/spinalcordtoolbox/PAM50; De Leener et al., 2017) it is now possible to investigate whether the signal-drop outs and their mitigation via z-shimming are also present in the gray matter (which is the relevant tissue for fMRI) and might even vary spatially (i.e. between dorsal and ventral horns). In order to address these two questions, we ran a 2×2 repeated-measures ANOVA (factor 1: condition with two levels: no z-shim, manual z-shim; factor 2: anatomical location: dorsal horn, ventral horn) where we tested for a main effect of condition, a main effect of location and an interaction between the two factors (Supplementary Material); this was followed up by post-hoc Bonferroni-corrected t-tests (where we also report % increase for the direct comparisons). As underlying metrics, we tested both the *mean* of signal intensity across all slices and the *variation* of signal intensity across slices. To assess robustness, the above-described analyses (based on single-volume EPIs) were repeated based on the average across the 250 motion-corrected volumes. As a negative control, we also performed the same analyses as above, but now splitting the spinal cord gray matter into left and right parts.

##### 2.6.1.5 z-shim effects on time-series data

The analyses described above, as well as the results reported by Finsterbusch et al. (2012) were solely based on measures of signal intensity. In order to directly investigate the potential benefit of z-shimming for spinal cord fMRI, we also investigated its effect on the temporal signal-to-noise ratio (tSNR, i.e. temporal mean divided by temporal standard deviation on a voxel-by-voxel basis) of motion-corrected data (250 volumes). We are aware that effects on tSNR do not allow for a perfect one-to-one extrapolation to effects on BOLD sensitivity, but we nevertheless believe this to be an adequate proxy measure due to the following reasoning (Deichmann et al., 2002; De Panfilis & Schwarzbauer, 2005; Poser et al., 2006): since the contrast-to-noise ratio (CNR) of BOLD responses is proportional to the product of the effective TE and tSNR and the effective TE does not depend on the magnetic field gradient in the z-direction, any tSNR gain obtained by z-shimming should reflect a corresponding relative gain in BOLD-CNR in arbitrary task-based fMRI studies.

Following up on section 2.6.1.4, we only assessed this in the region most relevant for fMRI, i.e. the gray matter of the spinal cord. We compared *mean* tSNR across all slices, as well as *variation* of tSNR across slices, between no z-shim and manual z-shim conditions: we descriptively reported % increase and also tested for significant differences using paired t-tests.

Since signal loss in the most caudal (inferior) slices in the no z-shimming condition could negatively impact the motion correction (as this is regularized along z using a 2^nd^-degree polynomial), we performed the above-mentioned analyses also after “censoring” of outlier volumes (Supplementary Material; see also section 2.5.1).

As we only acquired 25 volumes for the short and long TEs due to time constraints, we did not calculate TE-dependent z-shim effects on tSNR (as these would be based on unstable tSNR estimates).

#### 2.6.2 Automating slice-specific z-shimming

##### 2.6.2.1 EPI-based automation

Next, we investigated the performance of the EPI-based automated approach for selecting z-shim values, both in comparison to the conditions of no z-shim and manual z-shim; this was carried out in a sub-group of 24 participants. For the sake of brevity, we i) only reported our effects of interest – signal intensity based on single EPI volumes (Supplementary Material) and tSNR based on EPI time-series – in the spinal cord gray matter (i.e. ignoring whole-cord data) and ii) employed direct comparisons of conditions without using an initial omnibus test. Thus, in this sub-group of 24 participants we investigated: i) no z-shim vs manual z-shim, ii) no z-shim vs auto z-shim, and iii) manual z-shim vs auto z-shim. We reported % differences, as well as Bonferroni-corrected p-values from paired t-tests, again using *mean* and *variation* metrics.

##### 2.6.2.2 FM-based automation

We investigated the performance of the FM-based automated approach for selecting z-shim values (based on a different sub-group of 24 participants) using the identical procedure as outlined in the previous paragraph.

However, since we discovered that the performance of the FM-based approach was slightly inferior compared to the manual approach, we followed this up with several post-hoc investigations (detailed in the Supplementary Material). Briefly, we first used the vendor-based field map and assessed the contributions of i) the choice of mask for identifying the spinal cord in the field map phase data, ii) various choices of parameters employed in the fitting process of the gradient field, iii) field-gradients in the AP-direction, and iv) inhomogeneity-induced mis- localizations between EPIs and field map. Second, we substituted the vendor-based field map by the in-house field map and compared their performance. Third, we assessed the general reliability of estimating z-shim values from field map data by repeating the fitting process on a second in-house field map that was acquired at the end of the experiment. While all these attempts aimed to improve the estimation of the through-slice field inhomogeneity, a final modification of the approach involved a histogram-based evaluation of the observed field gradients in order to improve the resulting signal intensity.

##### 2.6.2.3 Comparing all three approaches

So far, the automated approaches were compared to the manual approach within each sub-group of 24 participants. We next turned to directly comparing the approaches, using all 48 participants.

First, we used two-sample t-tests (with Bonferroni-corrected two-tailed p-values) in order to assess the following, based on gray matter tSNR from EPI time-series (using both the mean as well as the variation of tSNR across all slices): i) comparing the baselines of no z-shim between the two groups, ii) comparing the improvement of manual z-shim vs no z-shim between the two groups, ii) comparing the improvement of auto z-shim vs no z-shim between the two groups and iv) comparing the difference of manual z-shim vs auto z-shim between the two groups. In complementary analyses, we also assessed the similarity between the automated approaches and the manual approach in terms of the actually chosen z-shim step using rank-based correlation and Euclidean distance (Supplementary Material).

Second, we assessed the stability of z-shim effects (based on either of the automated approaches as well as the manual approach) over time in all 48 participants. We were able to do this since we acquired an EPI reference-scan not only at the beginning of the experiment, but also at the end (∼60 minutes later). Using these reference scans, we ‘artificially reconstructed’ an EPI volume from each of the reference scans by selecting the corresponding volume for each slice based on the chosen z-shim values, no matter whether a participant was in the EPI-based or FM-based automation group. Importantly, we chose the ‘originally’ determined z-shim values to reconstruct ‘artificial volumes’ from both the first and the second reference scan. These volumes were then realigned to the mean of the motion-corrected time series. The warping fields that were obtained during the normalization of motion-corrected mean image to the template space were used to bring these volumes to the template space. We then compared gray matter signal characteristics (mean and variation of signal intensity across slices, respectively) for both time points using the various conditions via paired t-tests with Bonferroni correction.

### 2.7 Validation of EPI-based automation approach

In order to validate the EPI-based automation method (which performed at least as well as the manual approach), we obtained an independent, externally acquired data set of spinal GE-EPI data. These data were acquired by VO, RHD, and JCWB as part of a larger project on pharmacological aspects of cortico-spinal pain modulation (Oliva et al., 2022). Here, we report results based on analyzing the z-shim reference data from 117 acquisitions (39 participants, each with three visits).

The EPI reference scan (total acquisition time: 54 seconds) was acquired using a 2D EPI sequence with the following parameters: slice orientation: axial; slice number: 43 (20 slices for the spinal cord and 23 slices for the brain, i.e. concurrent cortico-spinal data acquisition); slice thickness: 4mm; slice gap: 25-50% (depending on the length of neck / size of head); field of view: 170×170mm^2^; in-plane resolution: 1.77×1.77mm^2^; TR: 3000ms; TE: 39ms; flip angle: 90°; GRAPPA acceleration factor: 2; z-shim resolution and range: 15 equidistant moments between -4.9 and 4.9 mT m^-1^ms (in steps of 0.7 mT m^-1^ms). The high-resolution T1-weighted images that were used for registration to template space were acquired with a 3D sagittal MPRAGE sequence with the following parameters: 260 sagittal slices; field-of-view: 320×260mm^2^; percentage phase field of view: 81.25%; voxel size: 1×1×1mm^3^; TE: 3.72ms; flip angle: 9°; TR: 2000ms; inversion time: 1000ms; GRAPPA acceleration factor: 3. All measurements were conducted on 3T whole body Siemens Skyra system.

As the validation dataset did not include volumes that were acquired under different z-shimming conditions, for each participant we ‘artificially reconstructed’ an EPI volume from their reference scan by selecting the corresponding volume for each slice based on the chosen z-shim values (see also section 2.6.2.3). We created three different EPI volumes for each participant and visit: i) a ‘no z-shim’ volume (based on an index of 8 for each slice, which corresponds to a z-shim moment of 0 mT m^-1^ms), ii) a ‘manual z-shim’ volume (based on the z-shim values manually selected by VO when the experiment was carried out) and iii) an ‘automated z-shim’ volume (based on the above-described EPI-based automation carried out post-hoc).

To bring these volumes to template space for each participant and visit, we applied the following steps to the T1-weighted anatomical data: i) segmenting the T1 image using SCT’s DeepSeg approach (Gros et al., 2019), ii) automatically labelling the vertebral levels C2-C7, and iii) bringing the T1 image to template space using non-rigid transformations. Then, we applied the following steps to the reconstructed EPI volumes: i) calculating the average of these three volumes (one volume for no z-shim, manual z-shim and automated z-shim each), ii) segmenting the average (using the PropSeg approach), iii) registering this average EPI to the template space (with the initial step of using the inverse warping field obtained from the registration of the T1-weighted image to the template image), iv) registering individual EPI volumes to the template space using the warps obtained from the previous step (in order to be unbiased), and v) in template space obtaining the signal over slices using the PAM50 cord mask.

Four individual data sets were excluded due to artifacts in the images (three data sets) and a wrong placement of the slice stack (one data set). Our final sample thus consisted of 113 measurements from 38 participants. Please also note that for preprocessing of data from one individual data set, we used a more recent version of SCT (version 5.2.0) due to a bug present in version 4.2.2.

Finally, we compared whole cord signal characteristics (*mean* and *variation* of signal intensity across slices) for i) no z-shim vs manual z-shim, ii) no z-shim vs auto z-shim, and iii) manual z-shim vs auto z-shim via paired t-tests with Bonferroni-correction and also reported % differences. For sake of simplicity, we treated each visit as a separate data point, thus ignoring the within-subject dependency structure. We also reported the results of the same analyses for gray matter signal characteristics (Supplementary Material).

### 2.8 Open science

The code that was run during the experiment for the automated selection of z-shim moments (both EPI-based and FM-based), as well as all the code necessary to reproduce the reported results, is publicly available on GitHub (https://github.com/eippertlab/zshim-spinalcord). Please also see the file Methods.md in this repository for a version of the Methods section with links to specific parts of the processing and analysis code. The underlying data are available in BIDS-format via OpenNeuro (https://openneuro.org/datasets/ds004068), with the exception of the external validation dataset obtained by VO, RHD and JCWB. The intended data-sharing via OpenNeuro was mentioned in the Informed Consent Form signed by the participants and approved by the ethics committee of the University of Leipzig.

Please note that during peer-review, the link to data will not yet work, as these will only be made public upon acceptance of the manuscript.

## Results

### 3.1 Replication and extension of previous findings

#### 3.1.1 Direct replication

Our first aim was to replicate earlier findings that demonstrated a significant increase of mean signal intensity and a decrease of signal intensity variation across slices via z-shimming. In our data set we were able to replicate these findings (Figure 2A), by also showing a significant increase of mean signal intensity (t_(47)_ = 19.97, p < .001, difference of 14.8%, CI: 13.4-16.2%) and a significant reduction of signal intensity variation across slices, either using the variance as a metric (as the to-be-replicated study did; t_(47)_ = 18.03, p < .001, difference of 67.8%, CI: 64-71.2%) or using the coefficient of variation (as we did in all further analyses; t_(47)_ = 23.97, p < .001, difference of 51%, CI: 47.7-53.8%).

**Figure 2.**
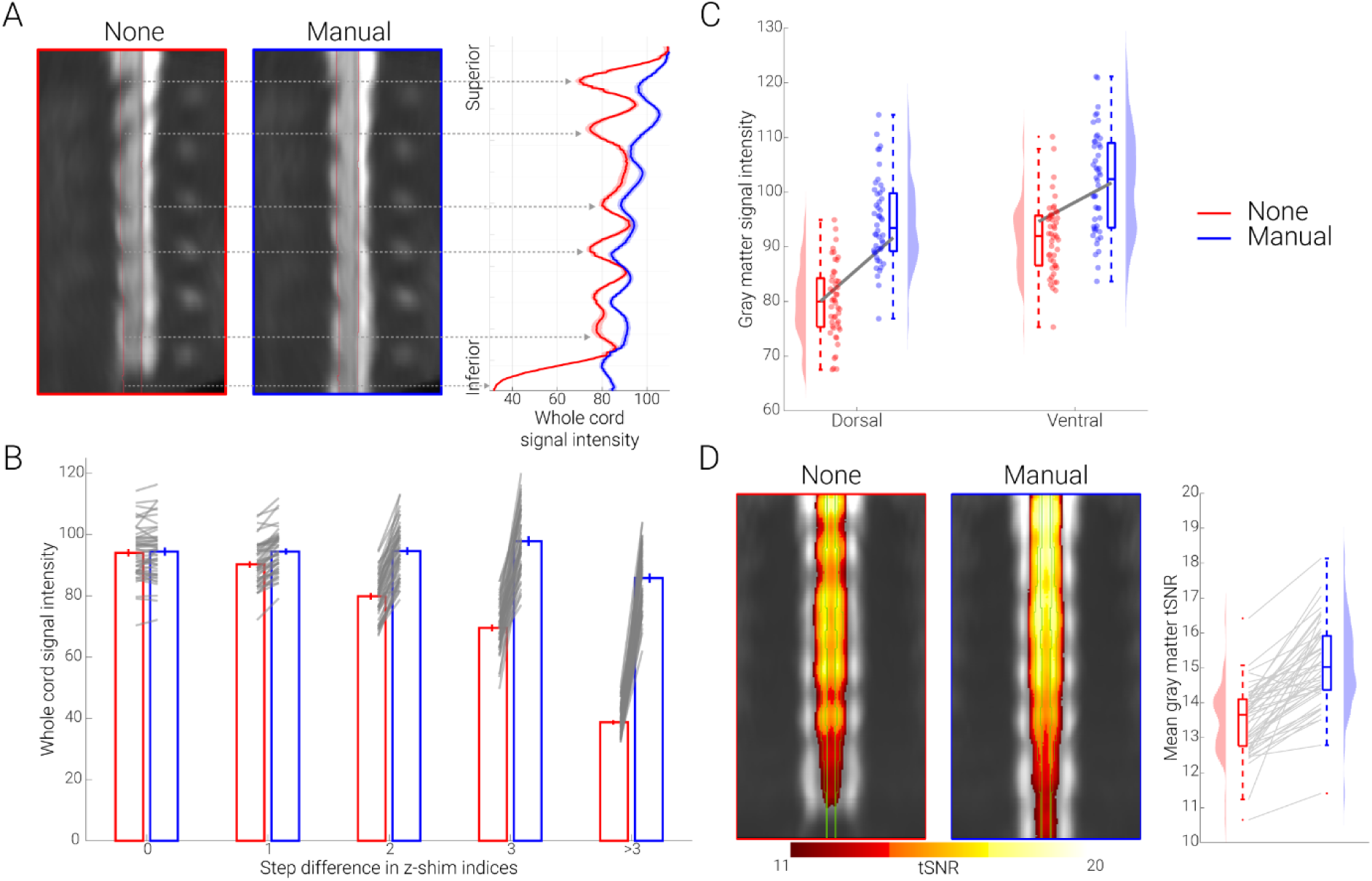
Replication and extension of previous results. A. Direct replication of Finsterbusch et al. (2012). The mid-sagittal EPI sections consist of the group-average single volume EPI data in template space of 48 participants acquired under different conditions (no z-shim and manual z- shim); red lines indicate the spinal cord outline. On the right side, group-averaged signal intensity in the spinal cord is shown for no (red) and manual (blue) z-shim sequences along the rostro-caudal axis of the cord. The solid line depicts the mean value and the shaded area depicts the standard error of the mean. **B. Slice-by-slice characterization of z-shim effects.** Bar graphs are grouped according to the absolute step size difference in the z-shim indices (x-axis) between no z-shim (red) and manual z-shim (blue) selections. The bars depict the mean signal intensity in the spinal cord for 48 participants for no and manual z-shim single volume acquisitions in native space. The vertical lines depict the standard error of the mean and the gray lines indicate participant-specific mean signal intensity changes between the no and manual z-shim conditions. **C**. **Z-shim effects in gray matter regions.** Signal intensity changes in different gray matter regions (dorsal horn, ventral horn) under different conditions (no z-shim, manual z-shim) are depicted via box-plots and raincloud plots. For the box plots, the median is denoted by the central mark and the bottom and top edges of the boxes represent the 25^th^ and 75^th^ percentiles, respectively, with the whiskers encompassing ∼99% of the data and outliers being represented by red dots. The circles represent individual participants and half-violin plots show the distribution of the gray matter intensity values across participants. The thick gray lines show the mean signal intensity across participants in the dorsal and ventral gray matter under different conditions. **D. Z-shim effects on time-series data.** Group-average coronal tSNR maps for the no z-shim and manual z-shim conditions as obtained from the motion-corrected EPI data in template space. The maps are overlaid onto the group-average mean image of the motion-corrected EPI data and depict a tSNR range from 11-20. The green line marks the outline of the gray matter. In the right panel, the participant-specific mean gray matter tSNR of the data acquired with and without z-shim are shown. Box plots are identical to those in C, the gray lines indicate individual tSNR changes between both conditions and the half-violin plots show the distribution across participants.

#### 3.1.2 Slice-by-slice characterization of z-shim effects

As depicted in Figure 2A, the improvement afforded by slice-specific z-shimming periodically varies along the rostro-caudal direction in a consistent manner across participants (for a depiction of individual data, see Supplementary Figure 1). In a next step, we thus investigated not only what the average benefit of z-shimming is across the entire slice-stack, but also quantified the benefit for slices with various degrees of signal-loss due to dropouts. We first asked what the maximal signal intensity gain is per participant and observed that this varied between 72% and 209%, with an average across participants of 122% (note that this analysis is based on the most-affected slice per participant). To descriptively characterize how many slices were affected by signal drop-out to what degree across participants, we quantified for each slice by how much the manually chosen z-shim value (between 1 and 21) differs from that of the no z-shim condition (a constant value of 11). We observed that on average 20% of slices had no difference, 32% of slices had a 1-step difference, 22% of slices had a 2-step difference, 11% of slices had a 3-step difference, and 16% of slices had more than a 3-step difference. In this last category, the most extreme possibly value (i.e. a 10-step difference) occurred only in 1% of the slices across the whole sample, demonstrating that the range chosen here for the z-shim reference scan is appropriate. As expected, signal intensity improvements became more pronounced with the increasing z-shim step size: 0% difference for a 0-step-difference, 5% different for a 1-step-difference, 18% difference for a 2- step-difference, 41% difference for a 3-step-difference and 122% difference for a >3-step- difference (Figure 2B); a statistical characterization of this relation can be found in the Supplementary Material.

#### 3.1.3 z-shim effects across different TEs

In addition to the TE of 40ms (which was the default across this study), we also investigated the effects of z-shimming at shorter (30ms) and longer (50ms) TEs. Focusing on mean signal intensity and signal intensity variation across slices, we observed a beneficial effect of z-shimming at the TE of 30ms (mean signal intensity: t_(47)_ = 18.82, p < .001, difference of 9.5%, CI: 8.6-10.5%; signal intensity variation across slices: t_(47)_ = 21.42, p < .001, difference of 48%, CI: 44.2-50.7% as well as at the TE of 50ms (mean signal intensity: t_(47)_ = 16.09, p < .001, difference of 11.6%, CI: 10.2-12.9%; signal intensity variation across slices: t_(47)_ = 22.20, p < .001, difference of 44.7%, CI: 41.4-47.7%).

#### 3.1.4 z-shim effects in gray matter regions

Next, we assessed whether z-shim effects might be present in the spinal cord gray matter and might even vary between the dorsal and ventral horns. An initially performed analysis of variance already indicated significant effects of z-shimming in the gray matter, as well as location-dependent effects of z-shimming (Figure 2C and Supplementary Material). Direct comparisons via Bonferroni-corrected paired t-tests revealed that there was a significant beneficial effect of z-shimming on mean signal intensity in the dorsal horn (t_(47)_ = 18.39, p < .001, difference of 18.2%, CI: 16.3- 20.3%), as well as in the ventral horn (t_(47)_ = 17.05, p < .001, difference of 10.9%, CI: 9.8-12.1%), but that the beneficial effect of z-shimming was more evident in the dorsal horn than in the ventral horn (t_(47)_ = 7.43, p < .001). These results are also in line with what can be observed visually in Figure 2A, where drop-outs seem to be most pronounced in the dorsal part of the cervical spinal cord (with the exception of caudal slices, where the whole cord is affected). As a negative control, we also performed the same analyses as above, but now splitting the spinal cord gray matter into left and right parts: as expected, there were no significant differences between these two regions.

#### 3.1.5 z-shim effects on time-series data

Moving away from reporting single-volume signal intensity measures, we next investigated the effect of z-shimming on the gray matter temporal signal-to-noise ratio (tSNR) of motion-corrected time-series data (250 volumes, acquired under no z-shim and manual z-shim, respectively). We observed a significant increase in mean tSNR (t_(47)_ =10.64, p < .001, difference of 11.9%, CI: 9.7- 14.2%), as well as a significant reduction of tSNR variation across slices (t_(47)_ = 11.01, p < .001, difference of 26%, CI: 21.9-30%), directly highlighting the benefits for spinal fMRI (Figure 2D). In the most-affected slices, z-shimming increased the tSNR by 28% on average, ranging from 1% to 155% across participants (this analysis also revealed that there was one outlier where tSNR decreased by 26% for manual z-shimming compared to no z-shimming).

### 3.2 Automation of z-shimming

The previous results were all obtained using manually determined z-shim values and we now turn to results obtained when automating the z-shim selection process, for which we propose two methods: one is based on obtaining these values from the EPI z-shim reference scan (EPI-based) and one relies on calculating the necessary z-shim values based on a field map (FM-based).

#### 3.2.1 EPI-based automation

In a sub-group of 24 participants, we first confirmed – using gray matter tSNR as obtained from motion-corrected time-series data – that also in this sub-sample manual z-shimming resulted in a significant increase in mean tSNR (t_(23)_ = 7.37, p < .001, difference of 10%, CI: 7.4-12.7%) and a significant decrease in tSNR variation across slices (t_(23)_ = 7.03, p < .001, difference of 27.2%, CI: 20.5-33.8%). Most importantly, we found a similarly beneficial effect when using our automated approach (Figure 3 upper panel; see also Supplementary Figure 3), i.e. a significant increase in mean tSNR (t_(23)_ = 8.69, p < .001, difference of 11.3%, CI: 8.9-13.9%) and a significant decrease in tSNR variation across slices (t_(23)_ = 7.04, p < .001, difference of 26%, CI: 19.4-32.7%). When directly comparing the two approaches to determine z-shim values, we observed no significant difference, neither for mean tSNR (t_(23)_ = 1.23, p = 0.70), nor for tSNR variation across slices (t_(23)_ = 0.61, p = 1), though a very slight benefit for the automated compared to the manual method was apparent.

**Figure 3.**
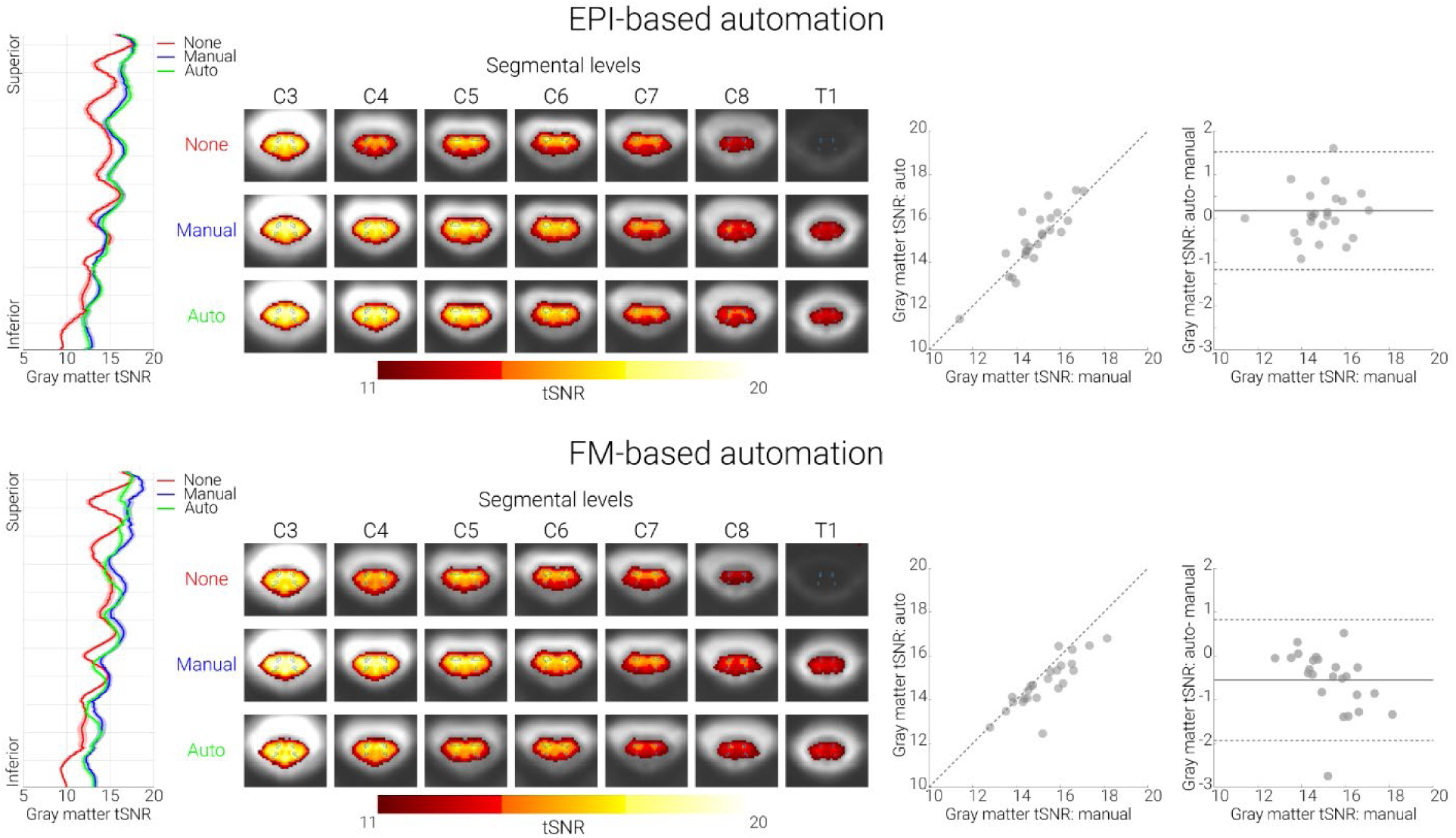
Performance of both automated methods. Top panel. EPI-based automation. **Bottom panel.** FM-based automation. In both panels, the left-most plots show the group- averaged gray-matter tSNR for no (red), manual (blue), and automated (green) z-shim sequences along the rostro-caudal axis of the cord. The solid line depicts the mean value and the shaded area depicts the standard error of the mean. Condition-wise group-average tSNR maps of the transversal slices at the middle of each segment are shown in the second graphs from the left. The maps are overlaid onto the group-average mean image of the motion-corrected EPI data and depict a tSNR range from 11-20. The outlines of the thresholded gray matter mask are marked by green lines. The scatter plots to the right show gray matter tSNR for manual (x-axis) and automated z-shim sequences (y-axis) plotted against each other (N = 24 for each automation sub-group). Bland- Altman plots show the gray matter tSNR for manual z-shim plotted as the ground truth (x-axis) and the difference in gray matter tSNR between automated and manual sequences plotted on the y-axis. The horizontal solid gray line represents the mean difference in the gray matter tSNR between the two (automated and manual) sequences, and the dotted lines show the 95% limits of agreement (1.96×standard deviation of the differences).

#### 3.2.2 FM-based automation

Field map data demonstrate that the source of the signal drop-outs z-shimming aims to compensate are B_0_ field inhomogeneities in the slice direction that i) are present where one would expect them based on anatomical and theoretical grounds (i.e. close to the intervertebral junctions and at the bottom of the field of view where the shim is poorer) and ii) are also consistent across participants (Supplementary Figure 2). In the FM-based approach, we therefore used field map data for z-shim calculation in a sub-group of 24 participants (different from the ones used for the EPI-based approach described above). We first confirmed – using gray matter tSNR as obtained from motion-corrected time-series data – that also in this sub-sample manual z-shimming resulted in a significant increase in mean tSNR (t_(23)_ = 7.99, p < .001, difference of 13.8%, CI: 10.6-17.4%) and a significant decrease in tSNR variation across slices (t_(23)_ = 9.36, p< .001, difference of 24.6%, CI: 20.4-29%). As expected, we also observed a beneficial effect of our FM-based approach, which resulted in a significant increase in mean tSNR (t_(23)_ = 6.41, p < .001, difference of 9.6%, CI: 6.9- 12.9%) and a significant decrease in tSNR variation across slices (t_(23)_ = 8.30, p < .001, difference of 21.8%, CI: 17.4-26.2%).

Unexpectedly though, despite this clear benefit, the performance of the FM-based approach was slightly worse than using manually determined z-shims (Figure 3 lower panel; see also Supplementary Figure 3): this occurred for mean tSNR (t_(23)_ = 3.86, p = .002), but not for tSNR variation across slices (t_(23)_ = 1.07, p = .88); please note that all p-values shown here and in the paragraph above are Bonferroni-corrected for three tests.

In post-hoc analyses carried out after the complete data-set was acquired, we investigated several possibilities that might account for this slightly poorer performance – all of these are explained in detail in the Supplemental Material. Briefly, we investigated the influence of i) the choice of mask for data extraction, ii) the choice of parameters for the fitting process, iii) the influence of field- gradients in the AP-direction, and iv) inhomogeneity-induced mis-localizations between EPIs and field map. We also investigated whether the type of field map played a role and whether z-shims could be reliably derived at all from field map data. These investigations aimed to improve the estimation of the through-slice field inhomogeneities in the field map. However, it should be noted that compensating the mean through-slice field inhomogeneity of a slice may not result in the optimum signal intensity: a few extreme values of the field inhomogeneity may shift the mean value significantly, thereby decreasing the signal of the majority of these voxels significantly; on the other hand, this shift may also not recover significant signal in the voxels with the extreme values, yielding an overall lower signal amplitude. To address this issue, a different approach of determining the z-shim value was used that was based on a histogram of the field gradients and aimed to reduce the influence of extreme values. This approach led to a consistent improvement in performance, although even this method still did not achieve the performance of the manual selection.

#### 3.2.3 Comparing all three approaches

To extend the within-group analyses reported above (each with N = 24) we next i) formally compared the three approaches based on the entire set of participants (N = 48) and ii) investigated the general question of how stable z-shim effects obtained via the three methods are across an experiment.

First, and most relevant for fMRI, we used mean gray matter tSNR to test for differences between the EPI-based and FM-based groups. These analyses (using Bonferroni corrected two-sample t-tests) revealed that there was neither a significant difference between the baselines of no z-shimming in the two groups (p = 1), nor a significant difference between the improvement compared to no z-shimming for either the manual (p = 0.38) or the automated approaches in the two groups (p = 1). However, we did observe a significant difference between manual z-shim vs auto z-shim in the two groups (p = 0.003), indicating the slightly worse performance of FM-based approach (see also Supplementary Figure 5). A second set of analyses based on tSNR variation across slices showed no significant differences between any of the approaches with all p-values > .9. The results of complementary analyses on how well the selected z-shim values matched between the manual approach and each of the automated approaches are reported in the Supplementary Material.

Second, we investigated how stable the beneficial effects of z-shimming are across time. When comparing how well each of the three z-shim methods performed against the case of no z-shimming in terms of mean signal intensity, we observed that despite some differences the beneficial effect of z-shimming was rather stable across time. More specifically, we observed that i) there was a significant difference between the two time-points in the baseline condition of no z-shim (t_(47)_ = 5.59, p < .001, with the first time point having significantly higher mean signal compared to second one), ii) that there was a slight degradation in performance when comparing z-shim benefits against no z-shimming between the 2^nd^ and the 1^st^ reference scan (manual: t_(47)_ = 8.44, p < .001; EPI-based: t_(47)_ = 9.70, p < .001; FM-based: t_(47)_ = 9.84, p < .001; thus similar across all three approaches) and iii) that all z-shim methods led to significant benefits not only in the data acquired at the beginning (manual vs no z-shimming: t_(47)_ = 22.35, p < .001, difference of 14%; EPI-based vs no z-shimming: t_(47)_ = 22.38, p < .001, difference of 14%; FM-based vs no z-shimming: t_(47)_ = 19.36, p < .001, difference of 11%) but also in the data acquired temporally later from when the z-shims were determined (manual vs no z-shimming: t_(47)_ = 18.52, p < .001, difference of 11%; EPI-based vs no z-shimming: t_(47)_ = 18.63, p < .001, difference of 11%; FM-based vs no z-shimming: t_(47)_ = 14.12, p < .001, difference of 8%).

### 3.3 Validation of EPI-based automation approach

In order to validate the EPI-based automation approach, we obtained an externally acquired corticospinal GE-EPI dataset consisting of 113 EPI z-shim reference scans acquired on a different MR-system (Oliva et al. 2022), which also allowed us to investigate the generalizability of the EPI-based automated approach in a dataset in which the manual selection was conducted by a different researcher (VO). Using this independently acquired data set, we observed that – compared to no z-shim – manual z-shimming resulted in a significant increase in mean signal intensity (t_(112)_ = 19.24, p < .001, difference of 22.1%, CI: 19.7-24.4%) and a significant decrease in signal intensity variation across slices (t_(112)_ = 8.83, p < .001, difference of 37.1%, CI: 29.7- 43.9%). Most importantly, the automated EPI-based approach resulted in a significant increase in mean signal intensity (t_(112)_ = 25.93, p < .001, difference of 28.3%, CI: 26.2-30.6%) and a significant decrease in signal intensity variation across slices (t_(112)_ = 10.98, p < .001, difference of 43.1%, CI: 36.4-49.3%). When we directly compared the automated and manual approaches, we observed that the automated method performed significantly better than the manual method both for mean signal intensity (t_(112)_ = 11.85, p < .001), and signal intensity variation across slices (t_(122)_ = 4.79, p < .001), demonstrating that the proposed EPI-based automated method can even outperform the manual selection (Figure 4).

**Figure 4.**
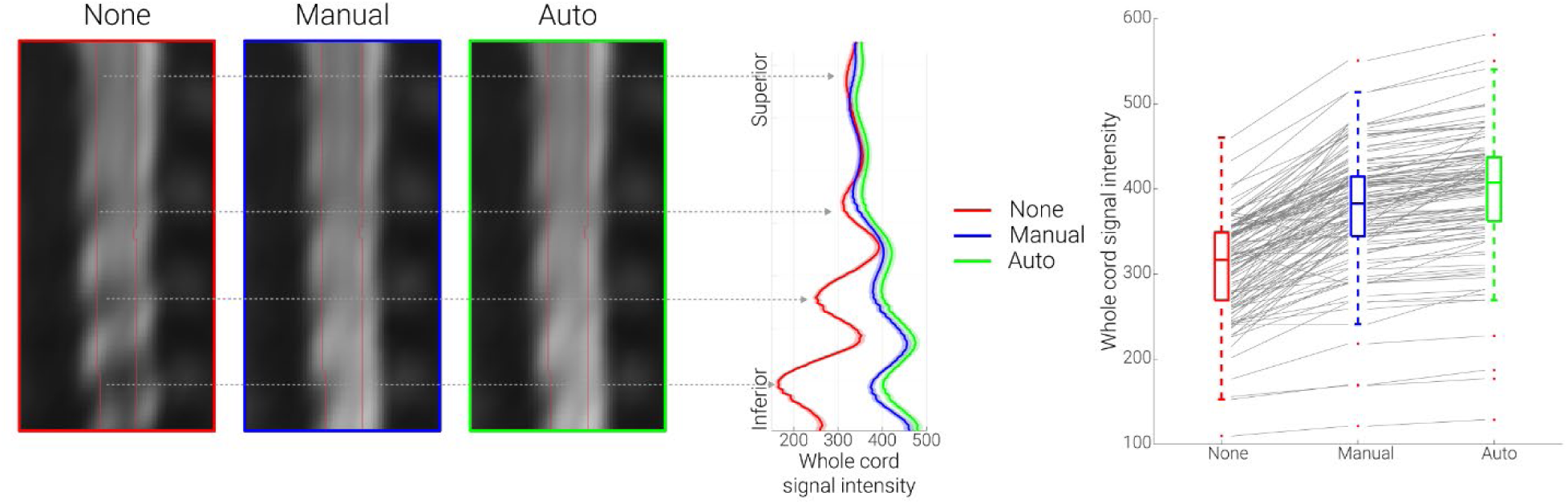
Validation of EPI-based automation on an independent data-set. The mid-sagittal EPI sections on the left consist of the group-average reconstructed z-shim reference scan EPI data in template space for the three different conditions (note that ‘EPI reconstruction’ was carried out via creating a single volume for each participant from the corresponding 15-volume z-shim reference scan by selecting for each slice the volume in which the z-shim moment maximized the signal intensity; for no z-shim reconstruction, the 8^th^ volume of the z-shim reference scan was selected, which corresponds to the central/neutral z-shim moment, as this acquisition had a range of 15 moments). The line plots in the middle depict the group-averaged spinal cord signal intensity (obtained from the reconstructed z-shim reference-scan EPIs) along the rostro-caudal axis of the cord for the different conditions. The solid lines depict the group-mean values and the shaded areas depict the standard error of the mean. The box plots on the right show the group-mean spinal cord signal intensity averaged over the entire slice-stack. The median values are denoted by the central marks and the bottom and top edges of the boxes represent the 25^th^ and 75^th^ percentiles, respectively. The whiskers encompass approximately 99% of the data and outliers are represented by red dots. The gray lines indicate the participant-specific data (N=113) upon which the box-plots are based.

## 4. Discussion

One of the main challenges in fMRI of the human spinal cord is the occurrence of spatially varying signal loss due to local magnetic field inhomogeneities. Here, we addressed this issue by employing the technique of slice-specific z-shimming. First, we aimed to replicate the results from the initial study on z-shimming in the spinal cord by investigating whether slice-specific z-shims mitigate signal loss in spinal cord GE-EPI data. Next, we probed the direct relevance of z-shimming to studies measuring spinal cord activity with fMRI, by investigating its benefits with respect to different TEs, gray-matter signals and EPI time-series metrics. Most importantly, we aimed to improve upon the typical implementation of slice-specific z-shimming (user-dependent shim selection) by developing two automated approaches: one based on data from an EPI reference-scan and one based on data from a field map acquisition.

### 4.1 Replication and extension of z-shim effects

The first demonstration of the benefits obtainable with slice-specific z-shimming in T2*-weighted imaging of the human spinal cord was provided by Finsterbusch et al. (2012), who developed a z-shim protocol tailored to the peculiarities of spinal cord imaging and assessed its effects on single volume GE-EPI data. Here, our first aim was to provide a direct replication of their results in a larger cohort of participants (N = 48) on a different MR-system. Similar to Finsterbusch and colleagues, we observed that z-shimming led to a significant and meaningful increase of average signal intensity (15%) and decrease of signal intensity variation over slices (68%) compared to the baseline of no z-shimming. In order to provide some detail on the expected benefits afforded by this method, we also performed a slice-by-slice characterization: while in ∼20% of the slices no z-shimming was needed, in the rest of the slices the application of a slice-specific z-shim resulted in a significant signal increase which could be as large as ∼200% in the most extreme cases. Comparing these effects to those obtained with slice-specific z-shimming in the brain (Deichmann et al. 2003; Weiskopf et al., 2006; Volz et al., 2019) – where z-shimming is critically important for signal recovery in susceptibility-prone regions such as the orbitofrontal cortex – it becomes clear that they are at least as prominent in the spinal cord and their compensation is thus critical in spinal cord fMRI.

The above-discussed results were obtained with a TE of 40ms in order to be close to the estimated T2* in the gray matter of the cervical spinal cord at 3T (41ms; Barry et al., 2019) and the TE considered by Finsterbusch and colleagues (44ms). Similar to Finsterbusch et al. (2012), we however also investigated the effect of z-shimming over different TEs (30ms, 40ms, 50ms), though now quantitatively and at the group-level. We observed that the beneficial effect of z-shimming was present to a similar degree across TEs, which is of direct relevance for fMRI. Longer TEs may be hard to avoid when covering lower cord sections due to the larger field of view required to avoid aliasing, in particular as higher in-plane acceleration factors may not be reasonable for the standard receive coils available. Conversely, shorter TEs might be desirable with respect to increasing the temporal resolution or optimizing BOLD sensitivity (Menon et al., 1993; Gati et al., 1997). In this respect, the consistent effect across TEs bodes well for using this technique flexibly in various settings.

In addition to the choice of TE in different scenarios of spinal fMRI, another important factor to consider is the anatomical region-of-interest. While this is typically the gray matter of the spinal cord – with studies on motor function likely focusing on the ventral horn and studies on somatosensation likely focusing on the dorsal horn – the specific effects of z-shimming on these structures are currently unclear, as Finsterbusch et al. (2012) only evaluated the entire spinal cord cross-section, thus averaging gray and white matter signals. There is indeed the possibility that z-shim effects might be rather negligible for the spinal cord gray matter, considering that field variations are most pronounced at the edge of the cord (Cooke 2004, Finsterbusch 2012, Cohen-Adad 2017), which largely consists of white matter. With the recent availability of probabilistic gray matter maps via SCT (De Leener et al., 2018), we were in a position to address this question in this study. We observed that the beneficial effects of z-shimming were highly significant and of appreciable magnitude in the gray matter. While these effects were already prominent in the ventral horns (11% increase), they were much stronger in the dorsal horns (18% increase) where signal losses were more severe (see also Cooke et al., 2004). Together, these results demonstrate the relevance of z-shimming for spinal fMRI and highlight its necessity specifically in studies of dorsal horn function, such as somatosensation and nociception. It should be mentioned though that it is currently unclear whether such a pattern will also hold outside of the cervical spinal cord, i.e. in thoracic and lumbar segments (see e.g. Finsterbusch, 2014). It is also important to note that in the current study, we aimed to optimize the signal in the entire spinal cord cross-section, but one might also consider optimizing the z-shim moments based on a gray matter region of interest. However, this approach would be more time-consuming (and might require user intervention), as for obtaining the gray matter masks it is necessary to first register the participant’s native-space data to template space and then warp the probabilistic gray-matter masks back to native space. Such a two-step approach is necessary since with the current spatial resolution and signal quality of EPI data at 3T it is not possible to automatically segment the gray matter robustly in every slice of every participant (in our experience, this also holds for T2*-weighted ME-GRE protocols in lower cervical segments).

The improvement in signal intensity we have discussed so far might in the worst case not directly translate into improved fMRI data quality (as indexed e.g. by tSNR): this might for example happen if physiological noise dominates the time-series or if participants move strongly in the z-direction and thus render the chosen z-shim moment for a slice incorrect. We therefore quantified the beneficial effect of z-shimming on gray matter tSNR, by acquiring time-series data under different z-shimming conditions, and observed a 12% increase in the mean tSNR and a 26% decrease of tSNR variability over slices. It is important to note that as none of the data analysed here were high-pass filtered or corrected for the presence of physiological noise (Brooks et al., 2008), it is likely that the absolute tSNR values observed (range across participants for manually z-shimmed data in template space: 11.4 to 18.1) represent a worst case. By acquiring z-shim reference scans at the beginning and at the end of our experiment (separated by ∼60 minutes), we were furthermore able to demonstrate that the effect of z-shimming was sufficiently stable across time, which is another important consideration for fMRI studies, as they usually require long scanning sessions. It should be mentioned however that participant-movement in the slice direction may reduce the performance of the z-shim compensation, although we deem this unlikely to happen frequently, considering the slice thickness used (5mm) and the distance of the vertebral disks that define the modulation of the magnetic flux density (∼15mm, see e.g. Wilke et al. 1997 for an overview and Busscher et al., 2010 for more recent data).

### 4.2 Automation of z-shimming

While the above-mentioned results demonstrate the utility of z-shimming for fMRI of the spinal cord, this approach requires detailed manual intervention in order to select the slice-specific z-shim moments. In order to overcome this drawback, in this study, we developed two different automated methods for the selection of the z-shim moments Although such approaches have been developed for fMRI of the brain (Weiskopf et al., 2007a; Marshall et al., 2009; Tang & Huang, 2011; Volz et al., 2019), they are lacking for fMRI of the spinal cord (with one notable exception to be discussed later, i.e. Islam et al., 2019) despite being desirable for a number of reasons. First, an automated method would be more time-efficient by reducing the time needed for selecting the z-shim moments. Second, it might enable more sites to perform spinal fMRI studies, as it reduces the need for extensive experience in judging spinal cord EPI data quality. Third, due to its automated nature it would eliminate the subjective (and error-prone) component involved in z-shim selection and thus increase reproducibility, which is especially important in longitudinal or multi-center studies. In the following, we describe the two different automated z-shim approaches we developed, with one being EPI-based and the other being field map based (FM-based).

The first automated method is based on the acquisition of an EPI z-shim reference scan – which is also employed for the manual selection – and relies on finding the z-shim moment that leads to the highest spinal cord EPI signal in each slice. This simple method achieved an at least identical performance in terms of all the investigated signal characteristics compared to the manual z-shim moment selection. In addition to that, the EPI-based approach was much faster compared to the manual selection: the calculations were completed in 15 seconds on average, whereas the manual selection took approximately 10 minutes for our set-up (24 slices and 21 z-shim values). The acquisition time of the z-shim reference scan was 55 seconds, but this could be shortened by limiting the range of the acquired z-shim moments. With the current set-up, we observed that the range of the acquired z-shim moments could indeed be restricted to achieve shorter acquisition times, as the most extreme z-shim moments were chosen quite rarely (lower-most moment of z-shim range chosen only in 1% of the slices, upper-most moment never). A drawback of the EPI-based approach is that it does not provide the flexibility to obtain slice-specific x- and y-shim settings at the same time in order to account for field gradients in the read and phase direction simultaneously (Volz et al., 2019). To obtain those, additional reference scans would be necessary and thus prolong the scan-time significantly (Finsterbusch et al., 2012). This drawback could be overcome by basing the slice-specific shim selection on a field map, which would allow for estimating x-, y-, and z-shims for each slice simultaneously – as already suggested by Finsterbusch et al. (2012) and Islam et al. (2019) – and this was the second approach we employed.

The FM-based approach was motivated by the fact that the source of the signal drop-outs which slice-specific z-shimming aims to compensate are B_0_ field inhomogeneities in the slice direction. The optimum z-shim value should thus be derivable from a field map, and we therefore fit a spatially linear gradient field in the slice direction to the measured field map data in order to estimate the gradient moment that will compensate the local through-slice field inhomogeneity. This FM-based approach provided highly significant benefits when compared to no z-shimming in terms of all the investigated signal characteristics. Similar to the EPI-based approach, the FM-based approach was clearly advantageous over manual z-shim selection in terms of the selection time (36 seconds on average). Compared to the EPI-based approach, it was however more time-consuming in terms of the time needed to acquire the different scans. First, we acquired a vendor-based field map, which took ∼5 minutes (though quicker field map acquisitions could be used). Second, we acquired a standard high-resolution T2-weighted image (which also took approximately 5 minutes to acquire; Cohen-Adad et al., 2021) for automated segmentation of the spinal cord, instead of the magnitude image from the field map. This choice was motivated by not wanting our results to be affected by the quality of the segmentation as neither the image resolution, nor the contrast of the magnitude image was optimal for the currently employed automated segmentation. However, the increased acquisition time for field map and T2-weighted image should be considered against the background that such images are acquired routinely in spinal fMRI experiments (e.g. for registration purposes). We thus believe that in typical research settings (where a few additional minutes of scan time might be negligible), the choice between the EPI-based and FM-based should in principle be guided by the slice-specific shim sets one needs to obtain (z-shim only: EPI-based; x-, y-, and z-shim: FM-based, though one would ideally want to acquire a field-map with higher resolution in the x-direction than done here).

However, we currently recommend using the automated EPI-based approach, as the performance of the automated FM-based approach was slightly inferior compared to the manual approach. While this difference was significant, it was small (∼4%) and limited to only some of the investigated metrics. We initially investigated several possibilities for this slightly worse performance (such as i) the choice of mask for data extraction, ii) the choice of parameters for the fitting process, iii) the influence of field-gradients in the AP-direction, iv) inhomogeneity-induced mis-localizations between EPIs and field map, and v) the reliability of FM-based z-shim calculation), but were not able to determine any factor that would improve the FM-based approach meaningfully. A slight but noticeable improvement was however brought about when substituting the vendor-based field map with a more robust in-house field map. A more significant improvement could be obtained by employing a histogram-based evaluation of the field gradients. Approximating the most probable field gradient values, this method aims to optimize the compensation for the majority of the voxels that contribute most to the signal, at the expense of more extreme values for which a significant compensation could only be achieved by sacrificing the compensation of most other voxels. While an improvement with this approach is observed, it still does not perform as well as the approach based on the reference scan which may have several reasons. On one hand, the relative intensities of the voxels as relevant in the EPI images are not appropriately considered. On the other hand, while the EPI-based approach is based on the same pulse sequence and has identical acquisition parameters as the target data (i.e. it exactly reflects the signal intensity achieved with the fMRI protocol), the FM-based approach is based on a different pulse sequence that is less prone to artifacts, but comes with a different voxel size as well as image orientation and position. These data could thus theoretically be expected to have a better quality and be more accurate, but may be less consistent with the EPI data (e.g. in terms of effects arising from in-plane field gradients (Deichmann et al., 2002; Weiskopf et al., 2007b) or slice thickness/profile modifications due to field inhomogeneities (Epstein & Magland, 2006) and most importantly are not determined from the EPI signal intensity.

It is also important to note that there are several ways of calculating the optimal z-shim moments from field map data and other approaches have for example taken the route of directly optimizing BOLD sensitivity in the brain based on EPI BOLD contrast models (e.g. Balteau et al., 2010; Volz et al., 2019). In the spinal cord, Islam et al. (2019) recently proposed an FM-based automated z-shim selection method for simultaneous brain and spinal fMRI. However, their implementation was aimed at compensating spatially broader field variations, as they fit a quadratic field term using voxels from slices that were ±4 cm distant from the target slice (which would cover 16 EPI slices in our case). In our study, we aimed to compensate for more localized field variations along the superior-inferior axis of the cord and therefore only included voxels from slices that were ±4 mm distant from the target slice. While comparing the performance of our approach directly to these approaches is beyond the scope of the current work, with the open availability of our code and data, this should be possible for the interested reader.

### 4.3 Validation of EPI-based z-shim automation

Finally, we demonstrated the validity of our EPI-based automation approach in an independently acquired large-scale cortico-spinal dataset (N > 100; Oliva et al., 2022). In this case, the automated approach exceeded the performance of manual selection (though we were not able to test this performance advantage in a further independent data-set). Such a pattern of results might be expected for studies where manual z-shim selection has to be performed rather fast due to time constraints (such as in the validation dataset, where a pharmacological challenge of the opioidergic and noradrenergic systems took place) – in our methodologically oriented study, particular emphasis was placed on the manual z-shim selection being as precise as possible, thus making the advantage of the automated approach possibly less apparent. This also hints at the potential of this approach to make z-shim selection more reliable and homogenous in complex studies where personnel might vary (e.g. in longitudinal or multi-site projects) and thus have different levels of experience that could detrimentally influence manual z-shim selection. Finally, since the cortico- spinal dataset naturally suffered from more severe signal drop-outs and acquisition artefacts such as ghosting (e.g. due to the large acquisition volume), the performance of the EPI-based automation approach demonstrates the robustness of this method with regards to varying levels of data quality.

### 4.4 Limitations

We would also like to point out several limitations of the presented work. First, the slice-wise z-shim approach is only applicable to axially acquired single-shot GE-EPI data. While this type of acquisition is used by numerous groups when studying somatomotor (e.g. Maieron et al., 2007; Vahdat et al., 2015; Weber et al., 2016; Kinany et al., 2019), somatosensory (Brooks et al. 2012; Tinnermann et al., 2017; Weber et al., 2020; Oliva et al., 2022) or resting-state spinal cord responses (Kong et al., 2014; San Emeterio Nateras, 2016; Kinany et al, 2020), there is also a strong tradition of using spin-echo approaches (for reviews, see e.g. Stroman, 2005 and Powers et al., 2018) and a more recent development in using multi-shot acquisitions (e.g. Barry et al., 2014; Barry et al., 2021; note that while the use of short TEs makes these acquisitions less affected by signal-dropout, in principle z-shimming might also be beneficial here). Second, although previous studies have demonstrated a high correlation of tSNR and signal intensity with BOLD sensitivity (particularly when effects of echo shifting are considered; e.g. Deichmann et al., 2003; Weiskopf et al., 2005; Poser et al., 2006), we cannot make direct extrapolations from the here-observed beneficial effects of z-shimming on tSNR to similar effects on task-based BOLD responses. In future methodological studies, it would thus be interesting to also acquire task-based spinal fMRI data under different z-shimming conditions to demonstrate the effect of z-shimming on the detection of BOLD responses – while this has been demonstrated in brain fMRI studies (Gu et al., 2002; Du et al., 2007), such evidence is currently lacking for the spinal cord (for a first step in this direction, see Islam et al., 2019). Third, the FM-based approach could be optimized e.g. by improving field-map quality to a degree where an automated segmentation of the magnitude image is possible (thus precluding any possible mismatch between the field-map and the T2-weighted image that is used for spinal cord identification) and increasing the spatial resolution of the field- map (currently limited at ∼2mm in x-direction) in order to allow for full xyz-shimming (see also Islam et al., 2019).

## 5. Conclusions

Spinal cord fMRI suffers from magnetic field inhomogeneities that negatively affect data quality, particularly via signal loss. In the current study, we extensively characterized the performance of slice-specific z-shimming in mitigating the effects of these inhomogeneities and developed two automated slice-specific z-shim approaches. We believe that our automated approaches will be beneficial for future spinal cord fMRI studies since they i) are less time-consuming than the traditional approach, ii) do not require extensive experience in judging data quality, and iii) are expected to increase reproducibility by eliminating the subjective component in the z-shim selection processes. This latter point is particularly important for longitudinal fMRI studies as they could be envisioned in the clinical setting where disease progression and treatment effects could be monitored.

## Acknowledgements

The authors would like to thank Torsten Schlumm for his programming work regarding the development of the in-house field map, the radiographers at MPI CBS for invaluable help with data acquisition and all volunteers for taking part in this study.

## Data availability statement

The underlying data are available in BIDS-format via OpenNeuro (https://openneuro.org/datasets/ds004068), with the exception of the external validation dataset obtained by VO, RHD, and JCWB. The intended data-sharing via OpenNeuro was mentioned in the Informed Consent Form signed by the participants and approved by the ethics committee of the University of Leipzig.

## Funding statement

JB received funding from the UK Medical Research Council (MR/N026969/1). FE received funding from the Max Planck Society and the European Research Council (under the European Union’s Horizon 2020 research and innovation programme; grant agreement No 758974). VO received funding from the Wellcome Trust (203963/Z/16/Z). NW received funding from the European Research Council under the European Union’s Seventh Framework Programme (FP7/2007-2013, ERC grant agreement No 616905), the European Union’s Horizon 2020 research and innovation programme (under the grant agreement No 681094) and the BMBF (01EW1711A & B) in the framework of ERA-NET NEURON.

## Conflict of interest disclosure

The Max Planck Institute for Human Cognitive and Brain Sciences has an institutional research agreement with Siemens Healthcare. Nikolaus Weiskopf holds a patent on acquisition of MRI data during spoiler gradients (US 10,401,453 B2). Nikolaus Weiskopf was a speaker at an event organized by Siemens Healthcare and was reimbursed for the travel expenses.

## Ethics approval statement

All participants provided written informed consent and the study was approved by the ethics committee at the Medical Faculty of the University of Leipzig.

## Supplementary Material

Please note that for sake of readability, the numbering of the headers in the Supplementary Material was kept consistent with the Results section in the main text.

### 3.1 Replication and extension of previous findings

#### 3.1.1 Direct replication

To additionally investigate how robust the findings in the main manuscript are, we supplement the single-volume analyses (that might be affected by various noise sources) by the same analysis approach, but now carried out on an EPI volume that is the average of a time-series of 250 motion corrected EPI volumes (acquired both for no z-shim and manual z-shim). We observed a significant increase of mean signal intensity (t_(47)_ = 19.03, p < .001, difference of 12%) and a significant reduction of signal intensity variation across slices (t_(47)_ = 27.22, p < .001, difference of 51%) for manual z-shim compared to no z-shim.

We also conducted the same analysis in native space (both for single-volume data and the average of a time-series of 250 motion corrected EPI volumes) instead of template space and observed very similar results demonstrating the benefit of z-shimming: for the single-volume data, we observed a significant increase of mean signal intensity (t_(47)_ = 21.07, p < .001, difference of 19%) and a significant reduction of signal intensity variation across slices (t_(47)_ = 25.55, p < .001, difference of 55%). For the average of a time-series of 250 motion corrected EPI volumes, we also observed a significant increase of mean signal intensity (t_(47)_ = 20.14, p < .001, difference of 16%) and a significant reduction of signal intensity variation across slices (t_(47)_ = 26.15, p < .001, difference of 54%). All of these results mirror those reported in the main manuscript.

#### 3.1.2 Slice-by-slice characterization of z-shim effects

Here, we complement the qualitative results reported in the main text by a more formal approach: we first carried out an analysis where we categorized each slice of the single-volume EPIs according to the step-difference between the manually chosen z-shim value and that of no z-shim (value 11) and compared the signal intensity in these categories between no z-shim and manual z-shim using a 2x5 repeated-measures ANOVA (factor 1: *condition* with levels no z-shim and manual z-shim; factor 2: *step-difference* of 0, 1, 2, 3, >3). We observed a significant main effect of condition (F_(1,88)_ = 222.74, p < .001), a significant main effect of step-difference (F_(4,352)_ = 355.8, p < .001) and a significant interaction (F_(4,352)_ = 204.66, p < .001). Post-hoc Bonferroni-corrected t-tests then revealed that the signal intensity improvement by z-shimming was not significant in those slices that had no step-difference and thus served as a negative control (t_(47)_ = 1.16, p = 1), but that it increased with increasing step-difference (step-difference of 1: t_(47)_ = 11.48 p < .001; step-difference of 2: t_(47)_ = 19.84, p < .001; step-difference of 3: t_(47)_ = 18.02, p < .001; step-difference of >3: t_(47)_ = 35.13, p < .001).

In order to assess the robustness of these effects, we also repeated the same analysis on the average of 250 motion corrected EPI volumes. The ANOVA showed a significant main effect of condition (F_(1,88)_ = 145.99, p < .001), a significant main effect of step-difference (F_(4,352)_ = 311.18, p < .001) and a significant interaction (F_(4,352)_ = 184.72, p < .001). Post-hoc Bonferroni-corrected t-tests revealed that the signal intensity in the no z-shimming condition unexpectedly was minimally (but consistently) higher in those slices that had no step-difference control (t_(47)_ = -4.95, p < .001, difference of 1%), but more importantly that the beneficial effect of z-shimming increased with increasing step-difference (step-difference of 1: t_(47)_ = 5.35 p < .001, difference of 3%; step- difference of 2: t_(47)_ = 16.91, p < .001, difference of 16%; step-difference of 3: t_(47)_ = 13.45, p < .001, difference of 32%; step-difference of >3: t_(47)_ = 32.79, p < .001, difference of 115%).

#### 3.1.3 z-shim effects across different TEs

When we repeated the analysis from the main text on the average of 25 motion-corrected volumes we observed very similar results. The effects of z-shimming were highly significant both at the TE of 30ms (mean signal intensity: t_(47)_ = 21.40, p < .001, difference of 10%; signal intensity variation across slices: t_(47)_ = 22.60, p < .001, difference of 48%) and at the TE of 50ms (mean signal intensity: t_(47)_ = 16.70, p < .001, difference of 12%; signal intensity variation across slices: t_(47)_ = 20.80, p < .001, difference of 44%).

#### 3.1.4 z-shim effects in gray matter regions

In order to formally compare the mean of signal intensity in different gray matter regions, we used a 2x2 repeated-measures ANOVA (factor 1: *condition* with levels no z-shim and manual z-shim; factor 2: *anatomical location* with levels dorsal horn and ventral horn). We observed a significant main effect of condition (F_(1,94)_ = 621.33, p < .001), a significant main effect of anatomical location (F_(1,94)_ = 39.70, p < .001) and a significant interaction (F_(1,94)_ = 21.31, p< .001); note that post-hoc t-tests are reported in the main text.

When we investigated the variation of signal intensity using the same ANOVA approach, we observed a significant main effect of condition (F_(1,94)_ = 1024.40, p < .001), a significant main effect of anatomical location (F_(1,94)_ = 30.32, p < .001) and a significant interaction (F_(1,94)_ = 9.60, p = .003). Following this up with post-hoc Bonferroni-corrected t-tests revealed there was a significant beneficial effect of z-shimming in the dorsal horn (t_(47)_ = 21.43, p < .001, difference of 48%), as well as in the ventral horn (t_(47)_ = 25.18, p < .001, difference of 47%), but that the beneficial effect of z-shimming was more evident in the dorsal horn than in the ventral horn (t_(47)_ = 4.06, p < .001).

As a negative control analysis, we carried out the same ANOVA approach, but now using the mean of signal intensity from the left vs right parts of the cord, where no differential effects should occur. As expected, we observed a significant main effect of condition (F_(1,94)_ = 690.05, p < .001), no significant main effect of location (F_(1,94)_ = 0.01, p = 0.90) and no significant interaction (F_(1,94)_ = 0.09, p = 0.76).

We repeated the above analyses (which are based on single-volume EPIs) with an average of the 250 motion-corrected volumes. A 2x2 repeated-measures ANOVA (factor 1: *condition* with levels no z-shim and manual z-shim; factor 2: *anatomical location* with levels dorsal horn and ventral horn) showed a significant a significant main effect of condition (F_(1,94)_ = 629.52, p < .001), a significant main effect of anatomical location (F_(1,94)_ = 36.55, p < .001) and a significant interaction (F_(1,94)_ = 4.90, p = .03). Post-hoc Bonferroni-corrected t-tests revealed there was a significant beneficial effect of z-shimming in terms of the signal intensity in the dorsal horn (t_(47)_ = 18.85, p < .001, difference of 15%), as well as in the ventral horn (t_(47)_ = 16.59, p < .001, difference of 11%), but that the beneficial effect of z-shimming was more evident in the dorsal horn than in the ventral horn (t_(47)_ = 4.87, p < .001). With respect to variation of signal intensity, the ANOVA resulted in a significant main effect of condition (F_(1,94)_ = 1300.20, p < .001), a significant main effect of anatomical location (F_(1,94)_ = 27.78, p < .001) and a significant interaction (F_(1,94)_ = 4.08, p = .046). Post-hoc Bonferroni-corrected t-tests revealed there was a significant beneficial effect of z-shimming in terms of reduction in the signal intensity variation over slices in the dorsal horn (t_(47)_ = 24.95, p < .001, difference of 46%), as well as in the ventral horn (t_(47)_ = 26.33, p < .001, difference of 48%), but that the beneficial effect of z-shimming was more evident in the dorsal horn than in the ventral horn (t_(47)_ = 2.69, p = 0.01).

#### 3.1.5 z-shim effects on time-series data

When we investigated the effects of z-shimming on tSNR using motion-censored time-series data, we observed a significant increase in mean tSNR (t_(47)_ = 10.73, p < .001, difference of 9%), as well as a significant reduction of tSNR variation across slices (t_(47)_ = 10.94, p < .001, difference of 25%). In the most-affected slices, z-shimming increased the tSNR by 26% on average, ranging from 6% to 116% across participants (this analysis revealed that there were 3 outliers where tSNR decreased by 1% (for two of the outliers) and 29% (for one of the outliers) for manual z-shimming compared to no z-shimming), again similar to what is reported in the main manuscript.

### 3.2 Automation of z-shimming

#### 3.2.1 EPI-based automation

When analyzing single-volume EPI gray matter signal intensity (in order to relate these effects to those from the direct replication performed earlier), we observed a significant increase of mean signal intensity (t_(23)_ = 12.51, p < .001, difference of 13%) and a significant decrease in signal intensity variation across slices (t_(23)_ = 16.89, p < .001, difference of 51%) for manual z-shimming against no z-shimming. Most importantly, we found a similarly beneficial effect when using our automated approach, i.e. a significant increase in mean signal intensity (t_(23)_ = 12.18, p < .001, difference of 14%) and a significant decrease in signal intensity variation across slices (t_(23)_ = 16.97, p < .001, difference of 48%). When directly comparing the two approaches to determine z-shim values, we observed no significant difference, neither for mean signal intensity (t_(23)_ = 0.31, p = 1), nor for signal variation across slices (t_(23)_ = 2.49, p = 0.06), though in both cases the performance of the automated approach was slightly superior. Overall, these results strongly mirror those based on tSNR reported in the main manuscript.

#### 3.2.2 Field map based (FM-based) automation

When analyzing single-volume EPI gray matter signal intensity, we observed a significant increase of mean signal intensity (t_(23)_ = 15.39, p < .001, difference of 15%) and a significant decrease in signal intensity variation across slices (t_(23)_ = 20.81, p < .001, difference of 52%) for manual z-shimming against no z-shimming. Most importantly, we found a similarly beneficial effect when using our automated approach, i.e. a significant increase in mean signal (t_(23)_ = 13.59, p < .001, difference of 12%) and a significant decrease in signal variation across slices (t_(23)_ = 17.42, p < .001, difference of 49%). When directly comparing the two approaches to determine z-shim values, we observed a significant difference for the mean signal intensity (t_(23)_ = 3.82, p = 0.003), but not for variation across slices (t_(23)_ = 1.52, p = 0.43), again showing the slightly inferior performance of this automated approach compared to manual z-shimming, congruent with the tSNR-based results in the main manuscript.

In the following, we detail the post-hoc investigations we undertook in order to determine possible reasons for the unexpected sub-optimal performance of the FM-based approach. Briefly, we first used the vendor-based field map and assessed the contributions of i) the choice of mask for identifying the spinal cord in the field map phase data, ii) various choices of parameters employed in the fitting process of the gradient field, iii) field-gradients in the AP-direction, and iv) inhomogeneity-induced mis-localizations between EPIs and field map. Second, we substituted the vendor-based field map by the more robust in-house field map and compared their performance. Third, we assessed the general reliability of estimating z-shim values from field map data by repeating the fitting process on a second in-house field map that was acquired at the end of the experiment. Finally, we calculated the optimum z-shim values using a histogram-based evaluation instead of a linear fit to reduce the influence of extreme values. In order to determine whether the performance of FM-based z-shim selection would improve with the different post-hoc approaches we undertook, we i) calculated the chosen z-shim values, ii) based on those we then artificially ‘reconstructed’ the EPI z-shim reference scan for each approach (see Methods section 2.7.3 for details), and iii) compared the gray matter signal characteristics (*mean* and *coefficient of variation*) between the new implementation and the original implementation. Please note that since we have a directional hypothesis (new FM-approach better than main manuscript FM-approach), we only test for an improvement compared to our original implementation.

##### 3.2.2.1 Choice of mask for identifying the spinal cord in the field map phase data

In our original implementation, the fitting of the linear gradient field was performed only on voxels within the spinal cord. This voxel selection was determined by a mask that was obtained from a segmentation of the T2-weighted image. While visual inspection of the mask overlaid onto the field map magnitude image did not give cause for concern in any of the 48 participants (i.e. due to possible participant movement between T2 and field map acquisitions), we nevertheless asked whether a change of the mask might improve performance. We therefore re-ran the original fitting procedure, but now based on a mask that was either eroded by 1 voxel or dilated by 1 voxel. When comparing the results based on these new masks to the standard mask, we observed that neither of these changes resulted in a meaningful and significant change in gray matter mean signal intensity (11% increase against no z-shim for all three masks) or signal intensity variation across slices (50% decrease for original and dilated masks, 49% decrease for eroded mask compared to no z-shimming). In line with these descriptive results, when directly comparing the original and new approaches statistically, we observed no significant differences (all p_uncorrected_ > 0.30).

##### 3.2.2.2 Choice of parameters employed in the fitting process of the gradient field

In our original implementation, we chose the following parameters based on pilot acquisitions: we smoothed the field map data with an isotropic 1mm kernel, used 9mm slab thickness (i.e. 9 transversal field map slices) for each fit and gave equal weight to all voxels in the fitting procedure. We next investigated whether variations of these parameters might have an influence on the performance: the smoothing kernel width (sigma) was set to 0, 1 or 2mm; the slab thickness was set to 5, 9 or 13mm, either with equal weighting or weighted by a raised cosine kernel of full-width half-maximum equal to the slab thickness and a roll-off factor beta of 0.5 (the purpose of the weighting was to down-weight voxels further away from the corresponding EPI slice). However, none of these choices seemed to make a meaningful difference, although out of these 17 additional variations (3 smoothing options crossed with 3 slab-thickness options and 2 weighting options) one showed a slight improvement for mean signal intensity and one showed a slight improvement for signal intensity variation along slices compared to our initial parameter set of choice that we used throughout the experiment (maximum improvement for mean signal intensity observed with parameter set “slab thickness = 9, beta = 0.5, smoothing sigma = 1”: t_(47)_ = 1.64, p_uncorreced_ = 0.054, p_corrected_ = 1, difference of 0.2%; maximum improvement for signal intensity variation over slices observed with parameter set “slab thickness = 9, beta = 0, smoothing sigma = 0”: t_(47)_ = 2.92, p_uncorrected_ = 0.003, p_corrected_ = 0.05, difference of 2%).Thus, the slightly worse performance of the FM-based approach reported in the main manuscript – which was based on a significant difference for mean signal intensity – does not seem to be due to the choice of parameters employed in the fitting process.

##### 3.2.2.3 Field gradients in the AP-direction

Another possible explanation for the slightly inferior performance of the FM-based approach is that field inhomogeneities in the A-P (y) direction may shift the center of k-space in the EPI acquisitions which – depending on their polarity – would result in a shorter or longer effective TE. Because the calculation of the required z-shim gradient moment from the field map assumes that the echo forms up at the nominal TE, any shift of the effective TE would lead to an imperfect compensation of the through-slice dephasing and would cause a signal loss. This is in contrast to the EPI-based approach, which rests on an EPI acquisition where the effective TE is inherently considered by just picking the best z-shim moment tested.

In the presence of a susceptibility-induced field gradient in the phase encoding direction, *G_SP_*, refocusing happens at an effective TE given by:

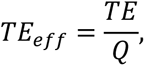

where

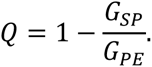

*G_PE_* is the effective phase encoding gradient:

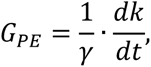

where *dt*is the echo spacing (Deichmann et al., 2002). Based on the fitted linear field gradient in the AP direction, we calculated *Q*for each slice, and adjusted the z-shim gradient moment to account for the effective TE. We then investigated how the adjustment of the z-shim moments (please note that we considered both positive and negative polarities of EPI phase-encoding) affected the gray matter signal characteristics compared to our original implementation. We neither observed a meaningful increase in mean signal intensity (negative polarity vs original implementation: t_(47)_ = -0.59, p_uncorreced_ = 0.72, p_corrected_ = 1, <0.1% decrease; positive polarity vs original implementation: t_(47)_ = 0.08, p_uncorreced_ = 0.47, p_corrected_ = 0.94, <0.1% increase) nor a meaningful decrease in signal intensity variation over slices (negative polarity vs original implementation: t_(47)_ = -0.93, p_uncorreced_ = 0.82, p_corrected_ = 1, 0.8% increase; positive polarity vs original implementation: t_(47)_ = -1.58, p_uncorreced_ = 0.06, p_corrected_ = 0.12, 1% decrease). It thus seems that the influence of AP gradients is rather negligible with respect to the slightly inferior performance of the FM-based approach.

##### 3.2.2.4 Inhomogeneity-induced mis-localizations between EPIs and field map

The FM-based z-shim selection relies on spatial congruency between the field map and the EPI acquisitions in the through-slice direction of the axial EPI volume. Local susceptibility-induced field offsets can however affect the spatial congruency in different ways. In the sagittal field map acquisitions, local field offsets will result in a shift along the readout direction, i.e. superior-inferior. The readout bandwidth of the field map acquisition was 630 Hz/pixel at a voxel resolution of 1 mm. Except for the region most inferior, the local field offsets were below 100 Hz, which results in voxel shifts of less than 0.16 mm. In the EPI-acquisitions, the field offset affects the effective slice localization. The EPI excitation bandwidth was ∼2 kHz at a 5 mm slice thickness. Field offsets <100 Hz would thus correspond to slice shifts <0.25 mm. In the worst-case scenario, where both effects are superimposed, an effective relative spatial shift of <0.4 mm is obtained which was deemed small enough to have a negligible impact on the z-shim selection.

However, to additionally empirically investigate whether an inhomogeneity-induced mis- localization between the EPI and the field map might be a driving factor for the slightly inferior performance of the FM-based approach, we selected the participants for which the FM-based automated selection of z-shim values led to a step-size difference of at least three steps in at least one slice compared to the manual z-shim values (N = 10). In other words, we tried to identify a sub-group with the most extreme differences, since higher step size differences compared to manual z-shim implies that the field map selection of z-shim values was unsuccessful or ‘off’. In those participants, we plotted the local field offset and the absolute difference between automated and manually selected z-shim values (Supplementary Figure 4) and then visually investigated whether there would be any detectable relationship between a high step size difference and high field offset. However, these plots do not indicate that higher step size differences generally coincide with high local field offsets.

##### 3.2.2.5 Use of different field map

We also investigated whether the quality of our default field map protocol might have led to the slightly inferior performance of the FM-based approach. In order to assess this, we calculated z-shim values not only based on the originally chosen field map (vendor-provided with 2 echoes), but also based on a separate in-house field map (with 12 echoes) which was acquired directly after the vendor-based one. When we quantified the signal characteristics, we observed that both methods led to a similar increase in mean signal intensity (11% for vendor-based and 12% for in-house field map) and decrease in signal intensity variation across slices (50% for vendor-based field map and 52% for in-house field map). In line with these descriptive results, when directly comparing the performance of the vendor-based and in-house approaches statistically (e.g. auto- vendor compared to auto-in-house; note that the baseline of no z-shimming is identical between the two), we observed a slight benefit of the in-house field map (for mean signal intensity: t_(47)_ = 2.34, p = 0.01, difference of 0.5%, for signal intensity variation over slices: t_(47)_ = 1.84, p = 0.04, difference of 4%). In order to test whether this improvement would lead to a change in the pattern reported in the main text (i.e. the FM-based approach performing worse when comparing mean tSNR for the automated compared to the manual approach), we followed up on this by comparing the performance of both approaches against manual approach and still observed a slightly inferior performance for FM-based approaches for the mean signal intensity (auto-vendor compared to manual: t_(47)_ = 7.14, p < .001, difference of 2.0%; auto-in-house compared to manual: t_(47)_ = 8.20, p < .001, difference of 1.5%) but not for the coefficient of variation (auto-vendor compared to manual: t_(47)_ = 0.46, p = 0.65; auto-in-house compared to manual: t_(47)_ = 1.40, p = .17), which is consistent with the results reported in the main text.

##### 3.2.2.6 Assessing the reliability of z-shim selection based on FM-based automation

To probe how reliably z-shims can in general be determined via field maps, we also acquired a second in-house field map near the end of our experiment (please note that due to technical problems the second field map was not acquired for three participants) and investigated whether this would result in similar automatically chosen z-shim values: across participants, we observed a mean Spearman rank-correlation of rs = 0.88, range: 0.50-0.98), suggesting that the robustness of the FM-based determination is unlikely to be a driving factor in the slightly inferior performance.

##### 3.2.2.7 Evaluating a histogram-based method of determining z-shims

In a further approach, we used a histogram-based method for automatically determining the slice-specific z-shim values from the field map data. This was based on the idea that for a broad distribution of field inhomogeneities, the chosen compensation gradient may only be able to recover significant signal for those voxels with a field inhomogeneity similar to that perfectly compensated. For a skewed distribution, the mean value may be shifted towards inhomogeneities that are less frequent which may reduce the overall signal accordingly. Thus, the chosen approach was based on the histogram of inhomogeneities and considered particularly the most frequent values. We first calculated the B_0_ z-gradient for all x- and y-values of each 1mm sub-slice of the vendor-based B_0_ map (swapped to the orientation of the EPI space) using the IDL procedure “gradient.pro” which, after proper scaling, resulted in a gradB_0,z_ map of the same resolution as the B_0_ map in mT/m. A histogram of gradB_0,z_ was then calculated for each EPI slice in a region-of-interest containing the spinal cord (as with all FM-based procedures, this was obtained from a segmentation of the T2-weighted image) of all of its five 1-mm sub-slices using a bin size of 0.01 mT/m. The resulting histogram was then smoothed with a kernel width of 1/20 of the total number of bins. Next, the main peak in the histogram was determined by comparing the surrounding of the three most frequent bins using the average of the respective center +/- 2 points. The final processing step for calculating a z-shim value for each EPI-slice was an weighted summation of the gradB_0,z_ bins within the range of the center +/- 10 points (corresponding to +/- 0.1 mT/m) around the resulting main peak with the constraint that the actual summation range was limited to points possessing more than 25% of the center’s intensity.

To demonstrate the improvement afforded by this method, we first show data from a single participant in which the original FM-based automation worked poorly in several slices. The left panel of Figure 6A shows that in an exemplary problematic slice, the z-gradient of the B_0_ map is not homogeneous across the spinal cord, leading to an asymmetric distribution of z-gradients and a sub-optimal choice of the z-shim value for this slice if the original approach is used (green line), differing also from the z-shim value determined by manual or EPI-based selection. Using the histogram-based method described above, the most probable value of gradB_0,z_ (gray line) is obtained by the intensity-weighted summation of the histogram around the main peak. As a consequence, the z-shim value obtained by this method better fits to that of the manual or EPI-based selection. In slices with more homogenous z-gradients across the spinal cord cross-section, both the original method and the histogram method provide virtually identical results (Figure 6A right panel).

We also assessed the improvement in signal quality offered by this method at the group-level, where we used the above-mentioned ‘reconstruction’ of the z-shim reference scan of each participant, using the slice-specific z-shim values suggested by the histogram-based method. Figure 6B shows that while this method did not completely eliminate the inferior performance of the FM-based approach, it led to a substantial improvement in signal quality across the group. When directly comparing the performance of the original FM-based approach to the histogram-based approach statistically, we observed a significant benefit of the histogram-based approach (mean signal intensity: t_(47)_ = 5.05, p < .001, difference of 1.3%; signal intensity variation over slices: t_(47)_ = 0.17, p = n.s.). We then followed up on this by comparing the performance of the histogram-based approach against the manual approach and still observed a slightly inferior performance for FM-based approach for the mean signal intensity (histogram-based compared to manual: t_(47)_ = 4.05, p < .001, a difference of 0.75%) but not for the coefficient of variation, in line with previous results. This minor penalty of the FM-based approach may be related to the fact that the relative signal intensities of the individual voxels as they contribute to the EPI image were not considered – this is in contrast to the approach based on the EPI reference scan. Together, this demonstrates that the evaluation of gradB_0,z_ by considering the corresponding histograms is capable of reducing the error in FM-based z-shim selection, even if it does not reach the performance of the manual approach.

#### 3.2.3 Comparing all three approaches

In order to compare how close the automated and manual (current ‘gold standard’) shim selection processes were, we calculated rank-based correlations and Euclidian distances between the chosen z-shim values in each of the two groups of 24 participants. This was done on a participant-by-participant basis for both metrics, which were based on the same input: slice-wise (i.e. 24) z-shim values from 1 to 21 (with 11 designating the neutral state of no z-shim) in the manual condition and in an automated condition. In both cases, we used two-sample t-tests to compare the two sub- groups (i.e. EPI-based automation and FM-based automation).

First, we calculated rank-based correlations between the values chosen for the manual and the automated approach. We observed very high correlations of z-shim values in the EPI-based group (average correlation: r_s_ = 0.95, t_(23)_ = 33.28, p < .001; range of correlations across participants: 0.85 - 0.99), as well as in the FM-based group (average correlation: r_s_ = 0.91, t_(23)_ = 26.04, p < .001; range of correlations across participants: 0.62 - 0.97). When directly comparing the two groups, we observed that the correlations were significantly higher in the EPI-based group (t_(46)_ = 2.67, p = .01).

Second (and overcoming the inherent limitations of a correlation-based approach, i.e. the fact that a perfect correlation might be obtained if the pattern of z-shim values were the same across slices, even if there was a constant shift in z-shim values), we employed the Euclidean distance – the square root of the sum of squared differences between the corresponding elements of the two vectors of z-shim values across slices – between the values chosen for the manual and the automated approach. We observed that while the average Euclidean distance for the EPI-based group was 3.40 (range across participants: 2.24–4.80), it was 5.21 for the FM-based group (range across participants: 3.61– 8.94), leading to a significant difference (t_(46)_ = 7.19, p < .001).

### 3.3 Validation of EPI-based automation approach

In the independently acquired data set, we observed that for gray matter signal intensity, manual z-shimming resulted in a significant increase in mean signal intensity (t_(112)_ = 22.04, p < .001, difference of 25%) and a significant decrease in signal intensity variation across slices (t_(112)_ = 8.29, p < .001, difference of 40%). When we investigated the performance of our automated EPI-based selection approach, we observed a significant increase in mean signal intensity (t_(112)_ = 28.44, p < .001, difference of 32%) and a significant decrease in signal intensity variation across slices (t_(112)_ = 10.32, p < .001, difference of 47%). When we directly compared the automated and manual approaches, we observed that the automated method outperformed the manual method both for mean signal intensity (t_(112)_ = 12.14, p < .001) and for signal intensity variation across slices (t_(112)_ = 5.63, p < .001).

## Supplementary Figures

**Supplementary Figure 1.**
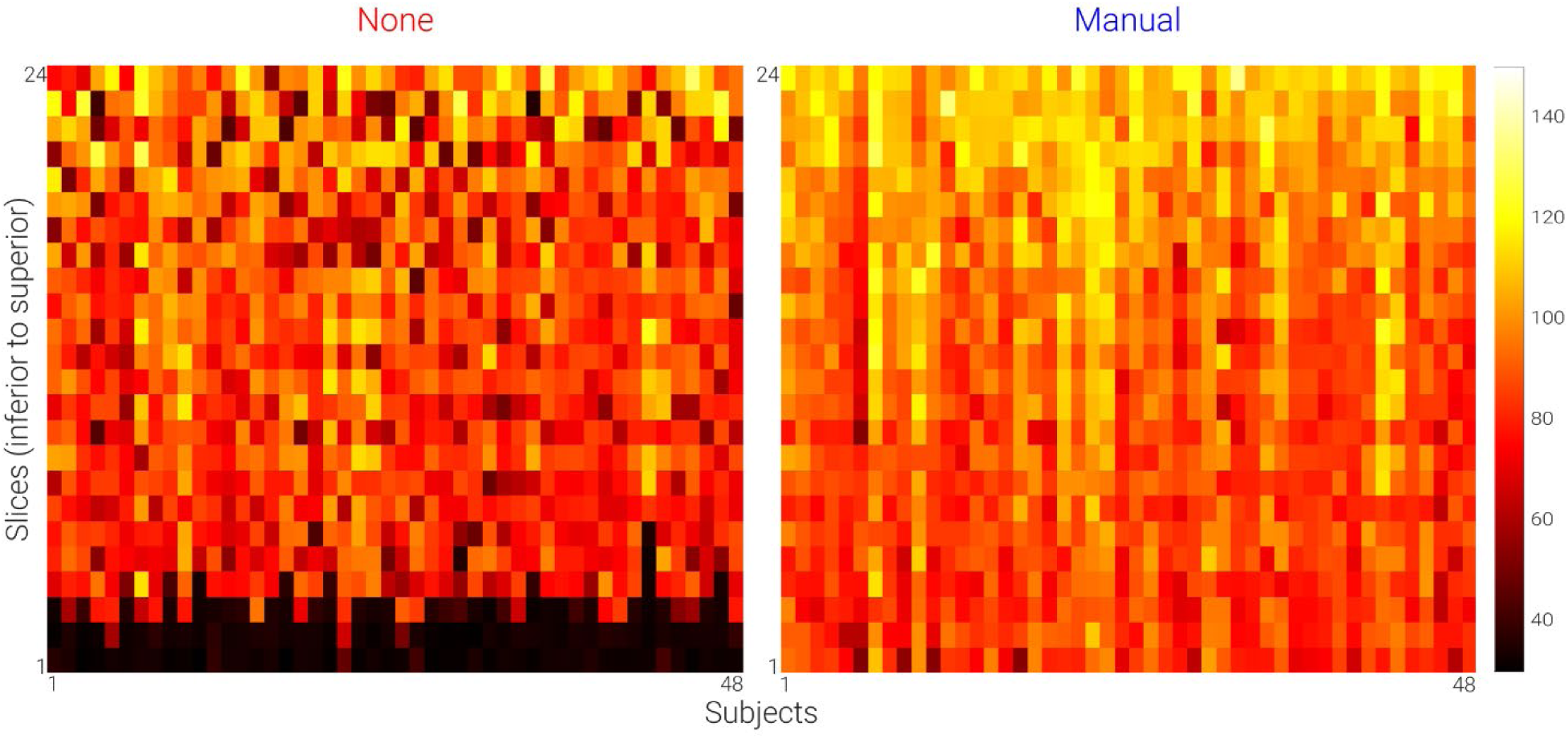
Slice-wise individual signal intensity data. Based on single volume EPIs acquired without z-shim and with manual z-shim, we calculated the mean signal intensity of each slice in native space. The heat-maps show signal intensity in axial slices (y-axis; 24 slices) for each participant (x-axis; 48 participants).

**Supplementary Figure 2.**
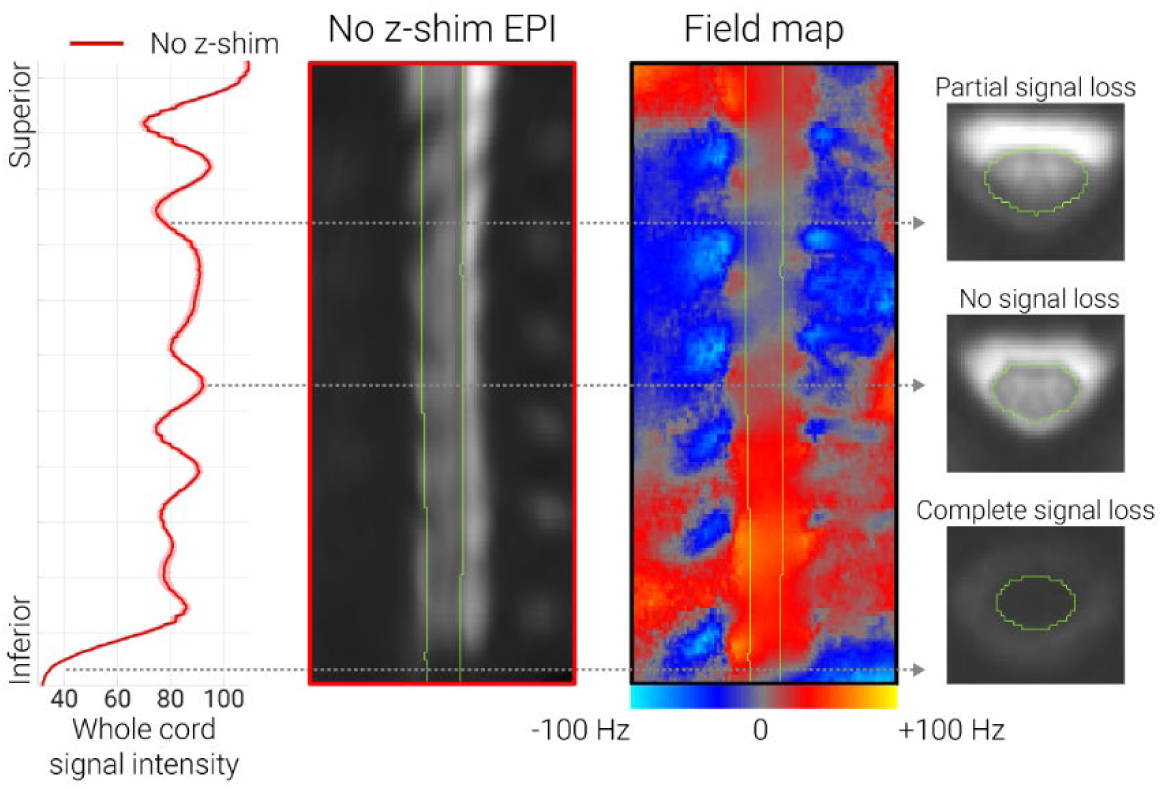
Relationship between field-variations and EPI signal loss. The line graph on the very left shows the group-averaged (N = 48) template-space spinal cord signal intensity along the rostro-caudal axis of the cord in acquisitions without z-shimming. The solid line depicts the group-mean value and the shaded area depicts the standard error of the mean. The mid-sagittal section on the left shows the group-average template-space single-volume EPI data acquired without z-shimming. The mid-sagittal section on the right shows the group-average template-space field map in order to depict the consistent field variations along the rostro-caudal axis of the cord. On the very right, there are three exemplary axial sections from the “no z-shim’ group-average template-space EPIs in order to demonstrate the influence of field variations on the EPI image quality in terms of signal loss.

**Supplementary Figure 3.**
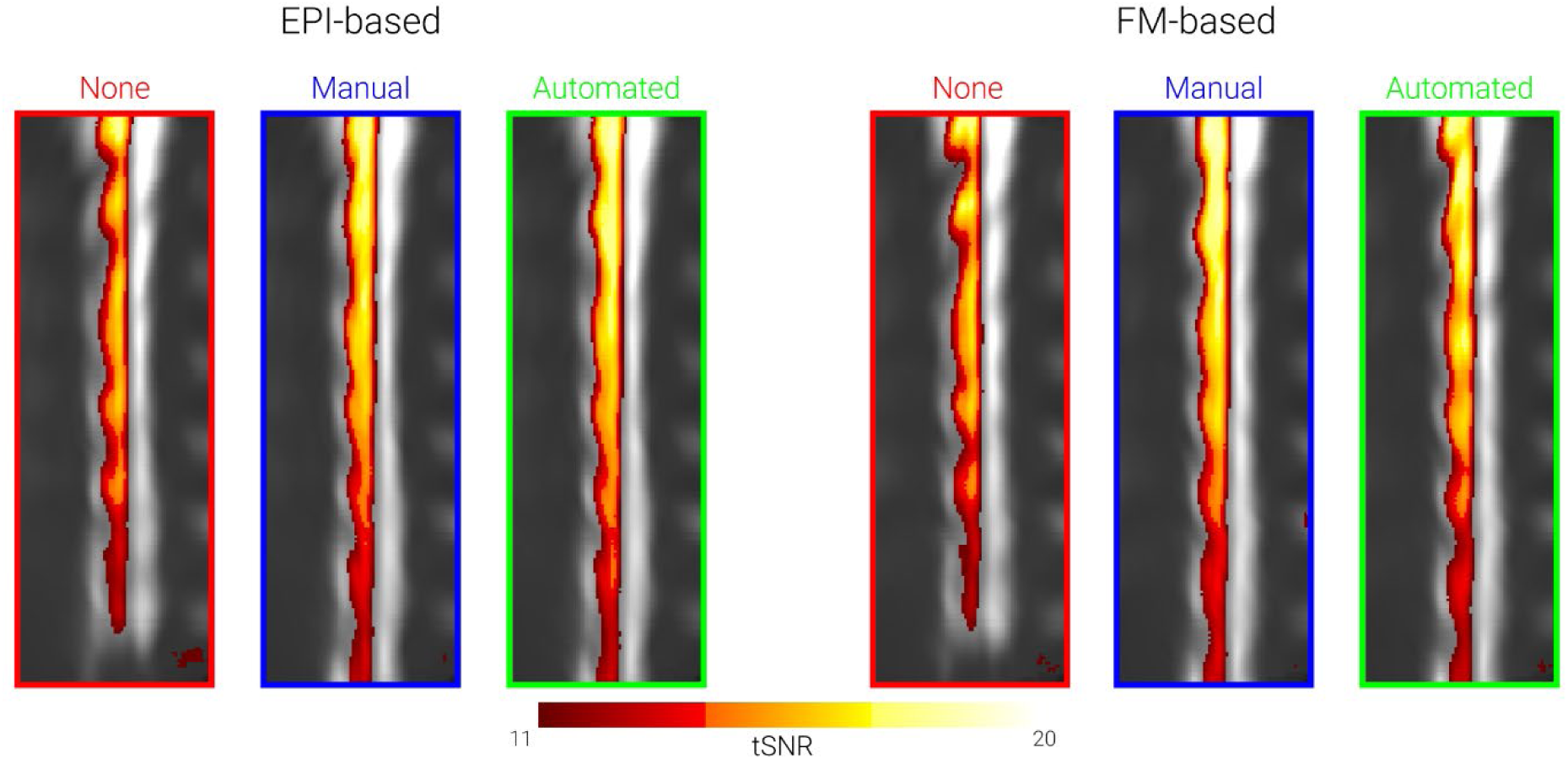
tSNR for different sequence variants. The mid-sagittal EPI sections in the background consist of the group-average mean of motion-corrected time-series data in template space for each sub-group of participants (EPI-based and FM-based, each of those with N=24) and condition (no z-shim, manual z-shim, automated z-shim). Condition-wise group-average tSNRmaps (based on the motion-corrected EPI data) are overlaid onto these mid-sagittal images (depicted tSNR range: 11-20).

**Supplementary Figure 4.**
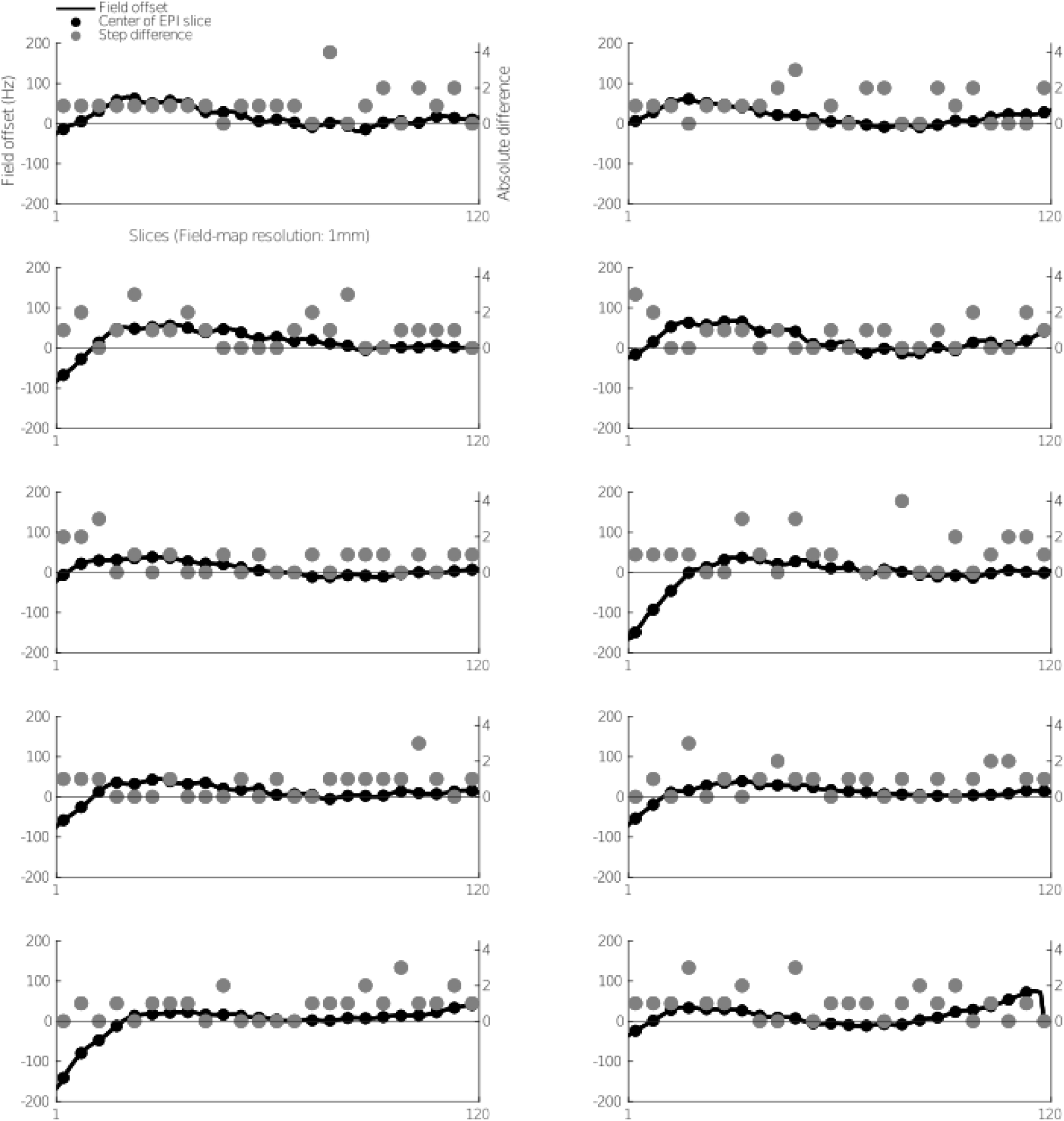
Relationship between field-offset and differential z-shim indices. Each subplot shows the field offset in Hz (black line; plotted on left y-axis) and the absolute difference in z-shim indices between the FM-based and the manual z-shim selection (gray circles; plotted on right y-axis). Depicted are those participants who had a difference of at least 3 steps between the FM-based and the manual z-shim selection (N = 10). Five FM slices (120 slices in total, 1mm slice thickness) correspond to a single EPI slice (24 slices in total, 5 mm slice thickness) with the black filled dots representing the corresponding center of each EPI slice in the FM resolution.

**Supplementary Figure 5.**
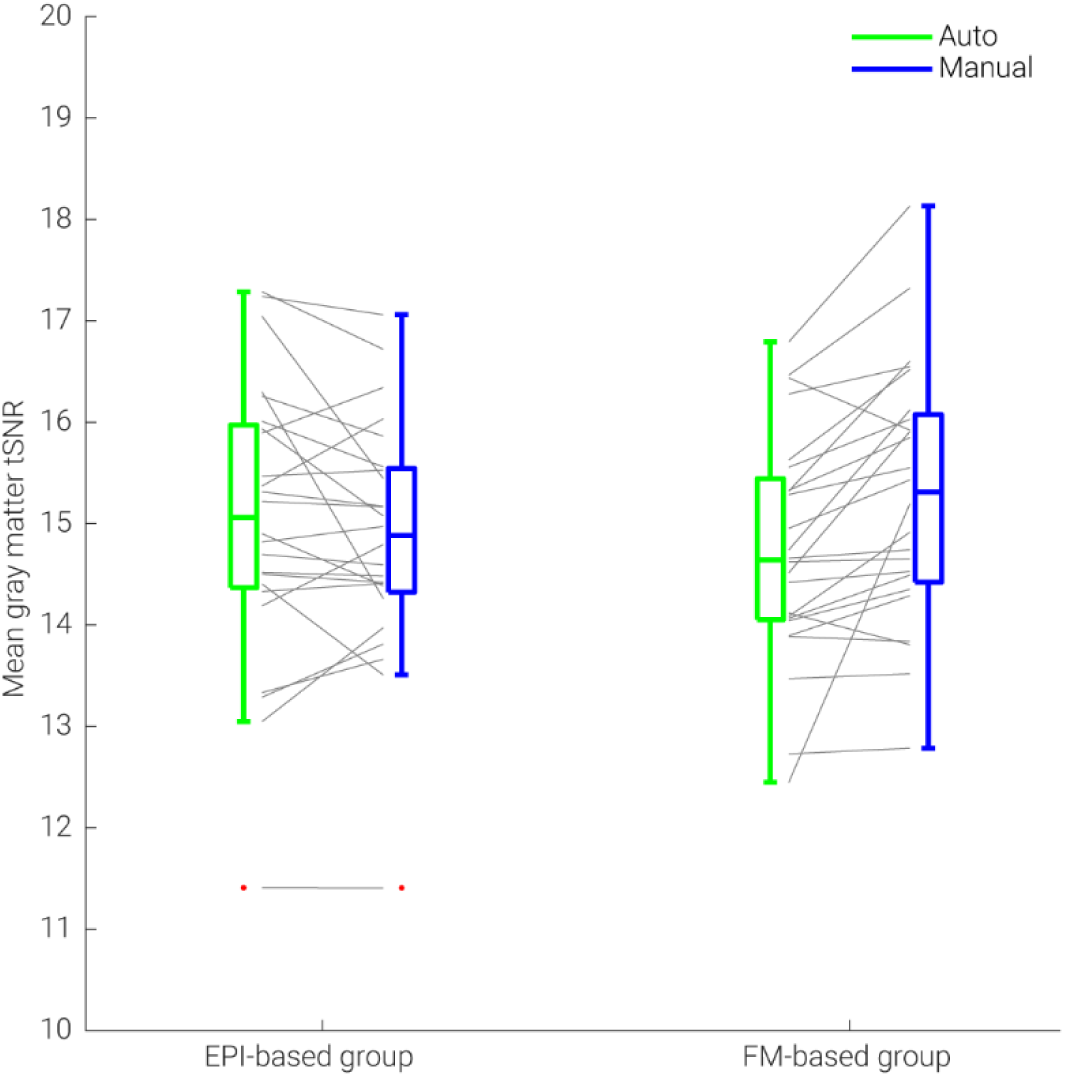
Mean gray matter tSNR of automated and manual approaches. We calculated the mean gray matter tSNR based on motion-corrected time series data acquired with different sequences (N = 24 for each group). The median is denoted by the central mark and the bottom and top edges of the boxes represent the 25^th^ and 75^th^ percentiles, respectively. The whiskers encompass approximately 99% of the data and outliers are represented by red dots. The gray lines indicate participant-specific tSNR in each condition and its change across conditions.

**Supplementary Figure 6.**
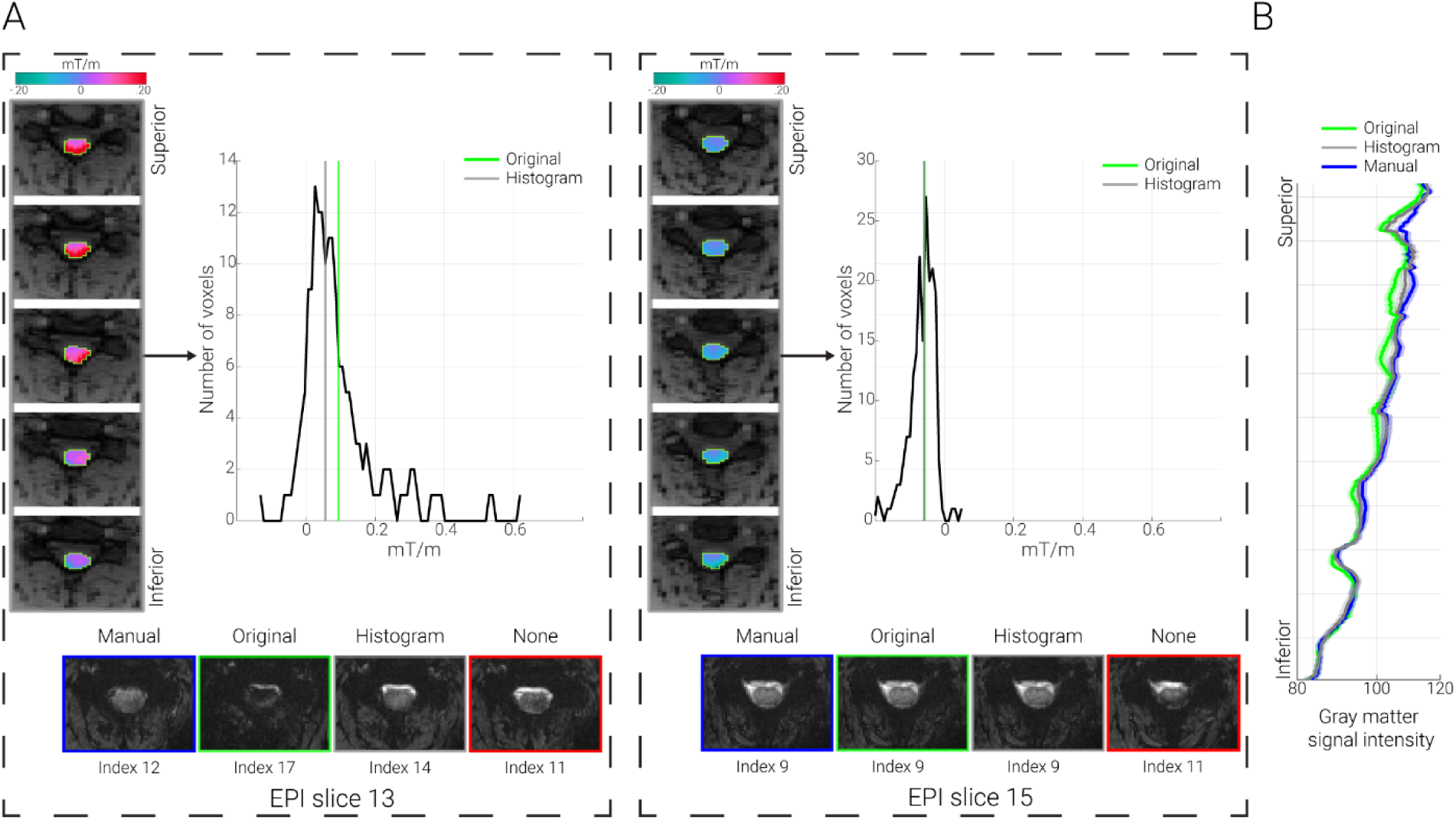
Histogram-based evaluation of field-map. A. Exemplary problematic & unproblematic slices. In both panels, five axial slices show the gradient map (gradB_0,z_) overlaid on the first magnitude image (for participant ZS030; in native space) corresponding to one EPI slice (problematic slice 13 and unproblematic slice 15, for left and right panels, respectively). The outlines of the cord mask (based on the T2-weighted image) are marked by green lines. The histograms show the gradB_0,z_ for these slices. On the lowermost part, the EPI volumes (corresponding to the selected z-shim indices) from the first z-shim reference image were taken for manual selection, original implementation, histogram-based implementation, and no z-shim condition for the relevant EPI slice. For slices with substantial field variation (problematic slice 13) the histogram-based shim offset selection offers clear improvement over the original automated approach. **B. Group-level signal intensity.** The line graph shows the group-averaged (N = 48) template-space spinal cord signal intensity along the rostro-caudal axis of the gray matter in the reconstructed EPIs (normalized) based on original FM-based implementation (green line), the manual selection (blue line), and based on histogram-based evaluation. The solid lines depict the group-mean value and the shaded areas depict the standard error of the mean.

